# Differential prevalence and risk factors for infection with coronaviruses in bats collected during 2020 in Yunnan Province, China

**DOI:** 10.1101/2024.05.28.596354

**Authors:** Ruiya Li, Alexander Tendu, Yakhouba Kane, Victor Omondi, Jiaxu Ying, Lingjing Mao, Shiman Xu, Rong Xu, Xing Chen, Yanhua Chen, Stéphane Descorps-Declère, Kathrina Mae Bienes, Meriem Fassatoui, Alice C. Hughes, Nicolas Berthet, Gary Wong

## Abstract

Coronaviruses (CoVs) pose a threat to human health globally, as highlighted by severe acute respiratory syndrome (SARS), Middle East respiratory syndrome (MERS) and the COVID-19 pandemic. Bats from the Greater Mekong Subregion (GMS) are an important natural reservoir for CoVs. Here we report the differential prevalence of CoVs in bats across biological and ecological factors within Yunnan Province. We also show the coexistence of CoVs in individual bats and identify an additional putative host for SARS-related CoV, with higher dispersal capacity than other known hosts. Notably, 11 SARS-related coronaviruses (SARSr-CoVs) were discovered in horseshoe bats and a Chinese water myotis bat by pan-CoV detection and Illumina sequencing. Our findings facilitate an understanding of the fundamental features of the distribution and circulation of CoVs in nature as well as zoonotic spillover risk in the One health framework.

## Introduction

Coronaviruses (CoVs) have gained global attention with the recent emergence of the COVID-19 pandemic, and prior to that, severe acute respiratory syndrome (SARS) and Middle East respiratory syndrome (MERS). However, whilst understanding the risk of spillover is necessary, a knowledge gap persists on the range of hosts for various types of CoVs. Such knowledge would provide a crucial foundation for future work to prevent the spillover of novel CoVs from either their natural hosts or susceptible intermediate hosts to humans.

Coronaviruses are classified into four genera: *Alphacoronavirus* (α-CoV), *Betacoronavirus* (β-CoV), *Gammacoronavirus* (γ-CoV) and *Deltacoronavirus* (δ-CoV), as defined by the International Committee on Taxonomy of Viruses (ICTV) in 2020 [1]. Some members of the β-CoV subgenus *Sarbecovirus* are known to cause severe disease in humans, including severe acute respiratory syndrome coronavirus (SARS-CoV) and severe acute respiratory syndrome coronavirus 2 (SARS-CoV-2), whereas severe acute respiratory syndrome-related coronaviruses (SARSr-CoVs) also pose potential risks. Over the past two decades, SARS-CoV, Middle East respiratory syndrome coronavirus (MERS-CoV) and SARS-CoV-2 have posed significant threats to public health [2, 3] with negative socioeconomic effects [4, 5]. Mitigating the adverse impacts of disease outbreaks caused by members of sarbecoviruses and merbecoviruses are a priority for public health authorities.

Coronaviruses infect a wide range of hosts: α-CoV and β-CoV can infect humans [6] and other mammals [7–10]. In addition, γ-CoV and δ-CoV have also been detected in wild birds [11] and poultry [12]. With notably different immune systems [13], bats are known to be important reservoir hosts for CoVs, especially as the hosts of the ancestors of five of the seven human CoVs, including human coronavirus 229E (HCoV-229E) and human coronavirus NL63 (HCoV-NL63) belonging to α-CoV, and MERS-CoV, SARS-CoV and SARS-CoV-2 belonging to β-CoV [14].

Understanding hotspots of potential vectors is crucial to understand the potential risk of spillover of novel CoVs. Rhinolophids appear to be a major reservoir of CoVs, and whilst distributed across the entire Old World, show the highest levels of richness in the Greater Mekong Subregion (GMS) [15] within Southeast Asia, though up to half of the rhinolophid species in the region are yet to be scientifically described [16]. The GMS includes territories from six countries: Myanmar, Thailand, Lao People’s Democratic Republic (Lao PDR), Vietnam, Cambodia, as well as Yunnan and Guangxi Provinces of China [17, 18]. Previous studies have shown that the GMS is a hotspot for a variety of emerging CoVs of bat-origin [19–25]. Over the past decade, two SARS-CoV-2 related coronaviruses (RshSTT182 and RshSTT200) were found in Shamel’s horseshoe bats (*Rhinolophus shameli*) collected from Cambodia in 2010 [19]. One SARS-related coronavirus (CoV) was identified in a Horsfield’s leaf-nosed bat (*Hipposideros larvatus*) collected from Eastern Thailand in 2013 [20]. RaTG13 was identified from an intermediate horseshoe bat (*Rhinolophus affinis*) sampled from Yunnan Province in 2013, and is understood to be among the closest relatives to the initial SARS-CoV-2 isolate responsible for the coronavirus 2019 (COVID-19) pandemic [21, 22]. Three new β-CoVs were detected in leaf-nosed bats (*Hipposideros*) sampled in Myanmar during 2016–2018 [23]. There is molecular and serological evidence indicating the presence of SARS-CoV-2 related coronaviruses (RmYN02 and RacCS203) from horseshoe bats (*Rhinolophus*) circulating in Southeast Asia in 2019 and 2020, respectively [24]. A SARS-CoV-2 related bat coronavirus (BANAL-236) was isolated in a Marshall’s horseshoe bat (*Rhinolophus marshalli*) captured in northern Laos in 2020, in which the viral spike protein was shown to mediate viral entry into hACE2-expressing human cells [25].

While these studies show the presence and diversity of SARS-CoV-2 like viruses among bat species endemic to the GMS, other CoVs are also present in the region. The scope of animal hosts susceptible to infection by these viruses, and their prevalence within the GMS, are not well characterized. To fill in these knowledge gaps, we identified CoVs in bats and determined their prevalence by host species and sampling locations across Yunnan Province, China. We then investigated the CoV diversity with focus on SARS-rCoVs in various bat species, with the view of further understanding the potential drivers behind the emergence of novel CoVs, their spread within animal populations and spillover to humans.

## Results

### Sample collection from various bat species from multiple trapping locations

A total of 716 bat rectal swabs (from 376 females/287 males, 53 no record), belonging to 17 bat species, were collected in 10 different caves (site A-J) of 3 locations (KM, YX and XSBN) (Figure 1-A, Table S1 and Table S4). The trapped bats belonged to six bat families including Pteropodidae (3 in total, 2 female /1 male), Hipposideridae (123, 75/46/2 unidentified sex), Vespertilionidae (228, 140/134/14), Miniopteridae (60, 31/20/9), Megadermatidae (2, both male), and Rhinolophidae (300, 160/103/37). Further details on sampled bats are shown in Table S1 and Table S4.

**Figure 1.**
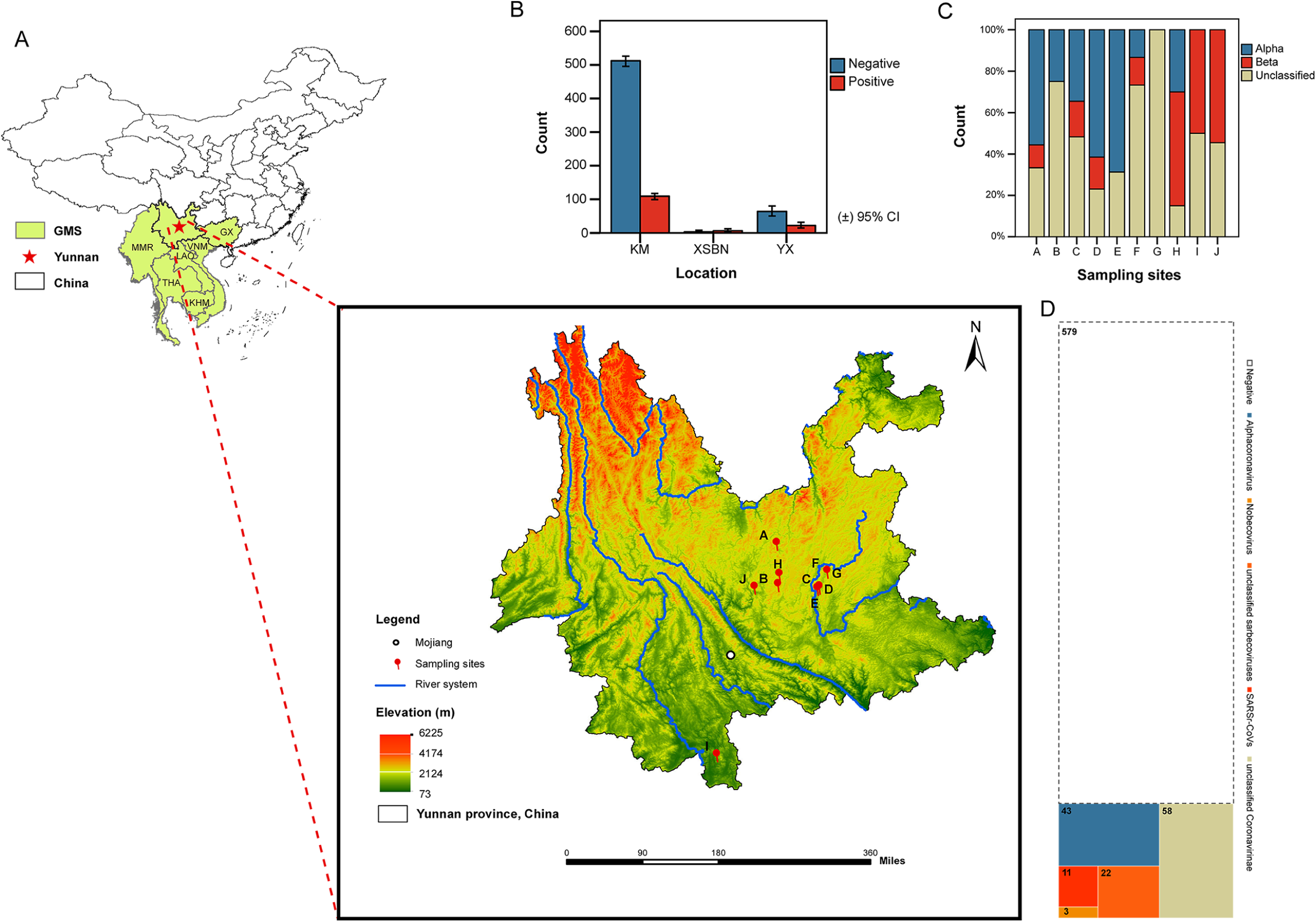
Geographical distribution of bat sampling sites and prevalence of coronavirus in the investigation. (A) The location of Yunnan province in relation to China and the Greater Mekong Subregion (GMS) and sampling sites. The white dot mark Mojiang county. Numbers and corresponding colors denote the different elevations (in Meters) in each place. A-J denotes ten different sampling sites from around Kunming (KM) City (sites: A-H), Yuxi (YX) City (site J) and Xishuangbanna (XSBN) Prefecture (site I) of Yunnan Province. Myanmar (MMR), Thailand (THA), Lao People’s Democratic Republic (Lao PDR), Vietnam (VNM), Cambodia (KHM), Guangxi Provinces of China (GX). (B) Coronavirus prevalence in three sampling locations (KM, XSBN and YX). (C) Coronavirus diversity in ten sampling sites (A-J). (D) Treemap in bat individual count for prevalence and diversity of coronavirus in investigation. Related to Table S1, Table S2 and Table S5.

### Coronavirus prevalence and correlation analysis of factors in rectal swabs from bats in Yunnan province, China

The overall CoV prevalence is 19.1% (95% confidence interval (CI): 16.2-22.0), which corresponds to 137 CoV-positive individuals (see Table S5 for details of BLASTn confirmation) out of 716 in total, and the negative rate is 80.9% (579 negative samples) (Table S1). The percentages and counts of each category are given for incidence by sites, species, genera and sex (Table S1 and Table S2).

#### Analysis of CoV prevalence based on sampling site

CoV prevalence was found to be associated with sampling site (Pearson χ^2^ (7, N = 699) = 29.791, p < 0.001) (Cramer’s V = 0.206, p < 0.001 and Goodman-Kruskal tau’s coefficient is 0.043, p < 0.001) (TableS3). Figure 2-C shows the pair-wise comparison of the CoV prevalence between pairs of sites. It was found that when the number of individuals exceeded 40 at each sampling site, CoV prevalence was between 5.8% (site B) and 32.6%* (site C) as shown in Table S1 and Figure 2B.

**Figure 2.**
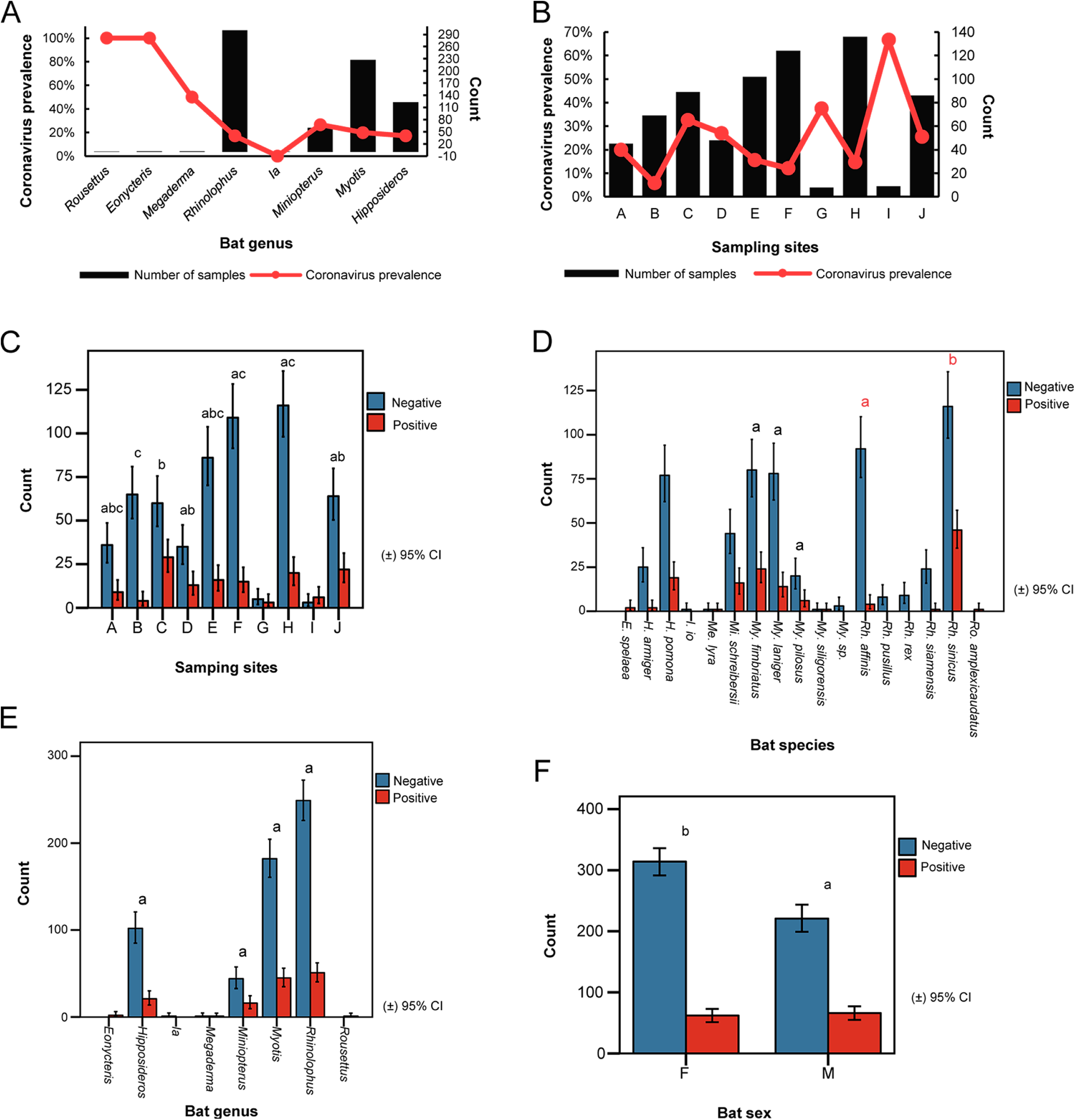
Coronavirus prevalence and analysis of significant correlation across different factors. (A) Prevalence of coronavirus and number of samples in each bat genus (B) Prevalence of coronavirus and number of samples in each sampling site. The x-axis indicates bat genera (A) and sampling sites (B). The y-axis indicates coronavirus prevalence (left) and the number of samples (right). The black column indicates the number of samples collected. The red line indicates coronavirus prevalence. (C) Coronavirus prevalence and significance of correlation with sampling sites. (D) Coronavirus prevalence and significance of correlation with bat species. (E) Coronavirus prevalence and significance of correlation with bat genera. (F) Coronavirus prevalence and significance of correlation with bat sex. Error bars represent 95% confidence interval (CI). Each small letter in the same colour above the bar (C-F) denotes a subset of categories whose column proportions do not differ significantly from each other (p≤0.05) by Chi-Square tests. Two small coloured boxes at the right corner indicate clusters of coronavirus positive and negative by different categories in the x-axis and the y-axis indicates the number of different categories in the two clusters accordingly in C-F. Related to Table S1, Table S2 and Table S6.

#### Analysis of CoV prevalence based on bat genera and species

Thirteen out of 17 species were found to be positive for CoVs as shown in Table S1 and Table S2. This was with the exception of *I. io, E. spelaea, Ro. amplexicaudatus* and *Me. lyra,* CoV prevalence was not associated with bat genus (Pearson χ^2^ (3, N = 710) = 3.473, p = 0.324 >0.05) (Figure 2-E) but was significantly associated with bat species within the genus *Rhinolophus* (Pearson χ^2^ (1, N = 258) = 22.648, p < 0.001). Here, *Rh. sinicus* were found to have a higher CoV prevalence than *Rh. affinis* (p < 0.05) (Figure 2-D). When the number of individuals sampled at a site exceeded 40 for a particular bat genus, CoV prevalence was between 17% (in *Rhinolophus* bats) and 26.7%* (in *Miniopterus* bats) (Figure 2-A). CoV prevalence across different bat genera and species are shown in Figure 2A, Figure 2D, Figure 2E, Table S1, and Table S7.

#### Analysis of CoV prevalence based on bat sex

CoV prevalence was 16.5% (95% CI: 12.7-20.3, 62 positives/376 samples) for female and 23.0% (95% CI: 18.1-27.9, 66/287) for male bats (Table S2). Further details are shown in Table S2 and Table S7. CoV prevalence was found to be associated with bat sex (Pearson χ^2^ (1, N = 663) = 4.424, p < 0.05) (Cramer’s V = 0.082, p < 0.05 and Goodman-Kruskal tau’s coefficient is 0.007, p < 0.05) (Table S6). For species where sample number exceeded ten, *Rh. sinicus* and *Rh. affinis* were found to have the highest and lowest CoV prevalence in both sexes respectively. When sample number exceeded ten in each genus, there was a trend of higher CoV prevalence in males rather than in female bats: *Hipposideros* (14.7% in females, 21.7% in males), *Miniopterus* (19.4%, 20.0%), *Myotis* (15.6%, 24.6%), *Rhinolophus* (16.9%, 20.4%) (Table S2 and Table S7).

### Differential prevalence of α-, β-, and unclassified-CoV is associated with sampling site, bat genus and bat species

All CoV positive samples contained a non-uniform composition of either α, β or unclassified CoVs (Table S3). α-CoV accounted for 31.4% (95% CI: 23.5-39.3, 43/137) of CoV positive samples (Table S3 and Table S5), whereas β-CoV accounted for 26.3% (95% CI: 18.8-33.7, 36/137) of CoV positive samples. The rest of the CoV positive samples (42.3%, 95% CI: 34.0-50.7, 58/716) belonged to unclassified CoVs.

Table S5 shows the best NCBI Blastn hit for each of the 137 CoV positive samples. We also show the phylogenetic analysis for all 137 positive samples in a polar tree (Figure 4-B). The compositions of CoVs (α-CoV, β-CoV and unclassified-CoV) in different bat species and sampling sites are shown in Table S3, Figure 1-C and Figure 3. Further deep sequencing (by MGI) confirmed the presence of α-CoV and unclassified-CoV in 9 CoV positive samples as shown in Table S8.

**Figure 3.**
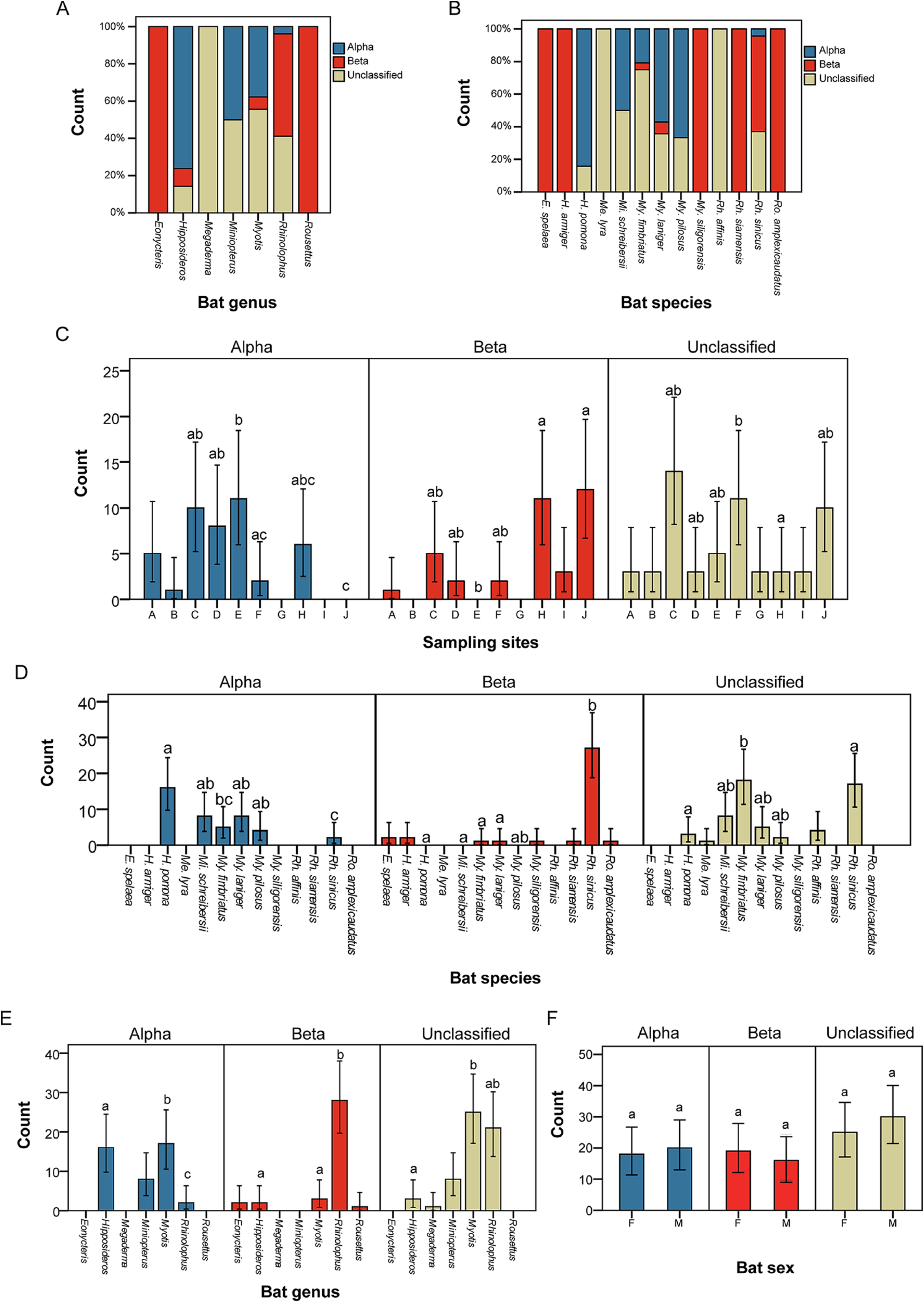
Differential prevalence of α-, β-, and unclassified-CoV and analysis of significant correlation across different factors. (A) Differential prevalence of CoV in 7 bat genra. (B) Coronavirus diversity in 13 bat species. The x-axis indicates bat genus (A) and bat species (B). The y-axis indicates the number of CoV-positives scaled to percentage. The yellow column indicates the number of unclassified coronavirus positives. The red column indicates the number of betacoronavirus positives. The blue column indicates the number of alphacoronavirus positives. (C) Differential prevalence of CoV in different genera and significance of correlation with sampling site. (D) Differential prevalence of CoV in different genera and significance of correlation with bat species. (E) Differential prevalence of CoV in different genera and significance of correlation with bat genera. (F) Differential prevalence of CoV in different genera and significance of correlation with bat sex. Each small letter above the columns of bar charts (C-F) denotes a subset of categories whose column proportions do not differ significantly from each other (p≤0.05) by Chi-Square tests. Each small colored box at the right corner denotes different categories in the x-axis indicating three clusters of α-, β-, and unclassified-CoV and the y-axis indicates the number of different categories in the three clusters accordingly in C-F. Related to Table S2, Table S3 and Table S9.

**Figure 4.**
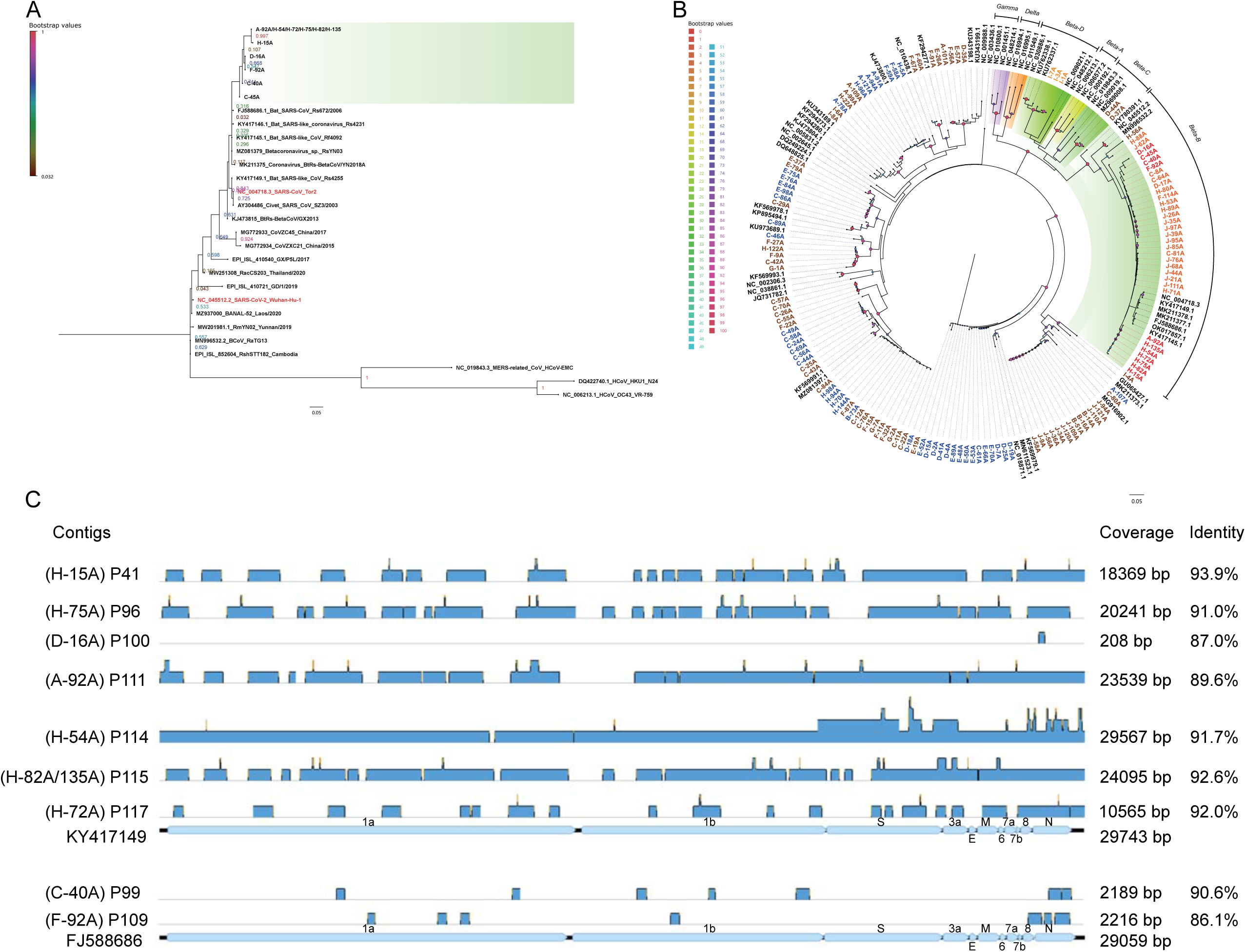
Phylogenetic analysis of targeted RdRp sequences of coronaviruses and identification of contigs in 11 SARSr-CoVs positive related libraries from NGS data. (A) Phylogenetic tree for the 11 SARSr-CoVs. The green block shows RdRp sequences for the SARSr-CoVs identified in this study while the entries in red are SARS-CoV and SARS-CoV-2. (B) Phylogenetic tree for all 137 CoV positives. (A) and (B) are both based on 195 bp targeted RdRp from Pan-CoV PCR. Seven sequences are less than 195bp, related to Figure S1. (C) The blue blocks represent contigs in NGS libraries corresponding to 11 SARSr-CoV positive individuals while mapped onto their best GenBank hits (KY417149 and FJ588686, related to Table S11). Phylogenetic analysis was conducted in MEGA11 using the Maximum-likelihood (ML) method, Tamura 3-parameter (TN92) model with Gamma distribution (*+G*) for Figure 4-A. General Time Reversible (GTR) model with Gamma distribution (*+G*) and evolutionarily invariable (*+I*) for Figure 4-B, which best model was selected in MEGA11 and the tree was established in QIAGEN CLC Genomics Workbench 22.0.2. The scale bar denotes the substitutions per site (0.05). Each letter number combination presents the targeted RdRp from bat rectal samples: alphacoronavirus is marked in blue, nobecoviruses are marked in light orange, sarbecoviruses are marked in red and unclassified sarbecoviruses are marked in orange, and unclassified coronavirus is marked in brown (based on the best hit from BLASTn). Others show reference sequences of coronaviruses. The different numbers below the branches indicate the support values of branch points. Node support was estimated from 1000 bootstrap replicates.

The differential prevalences of either α-, β-, or unclassified-CoV in bats was most significantly associated with sampling site (Pearson χ^2^ (10, N = 115) = 46.564, p < 0.001) (Cramer’s V = 0.450, p < 0.001 and Goodman-Kruskal tau’s coefficient = 0.197, p < 0.001) (TableS6). We also show pair-wise comparisons for differential CoV prevalence by site for pairs of sites (Figure 1-C, Figure 3-C and Table S3).

We show that differential CoV prevalence was also most significantly associated with bat genus (Pearson χ^2^ (4, N = 117) = 56.203, p < 0.001) (Cramer’s V = 0.490, p < 0.001 and Goodman-Kruskal tau’s coefficient = 0.223, p < 0.001) and bat species (Pearson χ^2^ (10, N = 125) = 81.907, p < 0.001) (Cramer’s V = 0.572, p < 0.001 and Goodman-Kruskal tau’s coefficient = 0.303) (Table S9). The data for the post-hoc pair-wise comparisons between bat genera and between bat species are shown in Figure 3-A, Figure 3-B, Figure 3-D and Figure 3-E.

Although CoVs showed a higher prevalence in male (23.0%) rather than female (16.5%) bats (Table S2 and Figure 2-F), we observed no significant difference in differential prevalence of α-, β-, and unclassified-CoV between male and female bats (Figure 3-F and Table S9). It was noted, however, that despite having a higher total CoV prevalence than all other male bats, male *Rh. sinicus* bats had a consistently lower prevalence of α-, β- and unclassified-CoV than female *Rh. sinicus* bats. Conversely, when the count of male and female both exceeded ten, male *H. pomona* (female: 14.3%/male: 21.05%) and *My. fimbriatus* (1.9%/8.2%) bats had a higher prevalence of α-CoV, whereas male *Rh. affinis* (0%/46.2%) had a higher prevalence of β-CoV. Among female bats, *H. armiger* (10.5%/0%) and *Rh. sinicus* (16.1%/0%) had the highest prevalence for β-CoV (Table S2).

### Beta-coronaviruses and SARSr-CoVs were detected in multiple genera of bats from Yunnan province

Of 36 β-CoV positive samples, 3 were *Nobecovirus* (“Beta-D” in Figure 4-B) and 33 were *Sarbecovirus* (“Beta-B” in Figure 4-D) positive, including 22 unclassified sarbecoviruses and 11 SARSr-CoVs (Figure 1-D and Table S4). The β-CoVs were detected in bats of the genera *Eonycteris* (2 samples), *Hipposideros* (2), *Myotis* (2), *Rhinolophus* (29) and *Rousettus* (1). The distribution according to bat species is shown in Table S5 and further genomic barcoding for bats found positive for SARSr-CoVs is shown in Table S10.

Following read assembly in NGS libraries sequenced from the 11 SARSr-CoVs positive individuals, contigs mapping to 35.5%-99.4% of the reference genomes (Pairwise identity: 89.6%-93.9%) for the corresponding SARSr-CoV references (GenBank references: KY417149.1 and FJ588686.1) were found in 9 corresponding NGS libraries. This was with the exception of library P100 (Individual ID: D-16A) from which a 208bp contig (Coverage: 0.7%, Pairwise identity: 87.0%) was obtained. For libraries P111 (A-92A), P114 (H-54A) and P115 (H-82A/135A), the contigs covered 11 major open reading frames encoding ORF1a, ORF1b, S, 3a, E, M, 6, 7a, 7b, 8 and N on the two references. Notably, over 99% of the whole genome was covered by the contigs assembled from library P114 (H-54A) (Figure 4-C and Table S11). Phylogenetic analysis showed that the 11 SARSr-CoVs may share a common ancestor with Bat SARS-CoV Rs672, Bat SARS-like CoV Rs 4231, Betacoronavirus RsYN03, Bat SARS-like CoV Rf4092 and BtRs-BetaCoV/YN2018 (Figure 4-A).

### Vertebrate-infecting viruses were also detected in SARSr-CoVs positive libraries

Six families of vertebrate-infecting viruses, including *Astroviridae*, *Circoviridae*, *Coronaviridae*, *Flaviviridae*, *Parvoviridae*, and *Polyomaviridae*, were detected in samples from SARSr-CoV positive individuals. In the ten libraries, most were rich in *Parvoviridae*. (Figure S2).

### Parasitized bats in northern Yunnan Province were found to be positive for α-CoVs

No obvious trend was observed between parasitized bats and CoV prevalence (Figure S3). The CoV-positive bats with ectoparasites were distributed across sites B, C, E, and H, all of which are distributed in the northern regions (i.e., the Kunming general area) of our sampling (Table S12). Among parasitized bats, α-CoVs were detected in three *My. pilosus* and three *Mi. schreibersii*, whereas the rest of the parasitized bats contained unclassified CoVs None of the β-CoV positive bats were parasitized (Table S10).

## Discussion

In determining the risk of CoV spillover, the surveillance of CoV prevalence in possible reservoir hosts is essential. Bats are an important order to study due to their demonstrated ability to host CoVs [26]. To investigate the occurrence of CoVs in bat populations, we determined CoV prevalence by pan-CoV semi-nested PCR and high-throughput sequencing, then proceeded to evaluate its correlation with risk factors across multiple sampling sites, different host genera, species, and sex in Yunnan Province, China. This region is of particular importance due to previous identification and/or isolation of human ACE2 utilizing relatives of human infecting β-CoVs [26–28].

### Habitat states (disturbance versus stability) and bat-specific factors influence the risk of virus spillover

Zoonotic spillover and transmission of emergent pathogens are influenced by both biological and ecological factors [29–34]. The impact of these drivers is poorly characterized but may contribute to varying ecophysiological stress among different bat hosts. They may include bat species, bat sex, life history traits, or ecological factors related to their habitats. While the biological factors e.g., sex, and species are immutable, habitat quality may be disturbed. This further exacerbates physiological stress in these hosts, a factor that has been shown to increase pathogen spillover risk [35, 36]. For these reasons, location, bat species and bat sex are important and fundamental considerations in identifying risk factors that contribute to the initial spillover and spread of bat-borne CoVs to new hosts [35, 37].

### CoV prevalence and is associated with sampling site

In this investigation, the prevalence of bat-borne CoVs ranged between 5.8-66.7% for all ten sampling sites (A-J) (Figure 2-B, Figure 2-C and Table S1) and between 17.6%- 66.7% for the three categories of sites (KM, YX and XSBN) (Figure 1-B and Table S1). Although statistically significant, the coefficient of correlation was low, indicating a weak correlation between site and CoV prevalence in Yunnan. For sites with greater than 40 bats, CoV prevalence was found to be highest in site C in Kunming (Figure 2-B), with significant differences between sampling sites as shown in Figure 2-C.

The differential prevalence of CoVs, α-CoV, β-CoV and unclassified CoVs were significantly associated with sampling sites, bat genera and species (p < 0.001) (Table S6). Kunming and Yuxi, in northern Yunnan, differ substantially in climate and altitude from Xishuangbanna prefecture (XSBN) in southern Yunnan (Figure 1). We noted a statistically significant difference in β-CoV prevalence between the northern (Kunming and Yuxi) and southern (XSBN) sites, with XSBN showing a higher β-CoV than Kunming and Yuxi combined (Table S3). Greater sampling in these regions would illuminate on whether this is merely the result of a small sample size in XSBN, considering that the highest proportion of β-CoV positive bats were found in Yuxi.

### *My. laniger* may host relatives of β-CoVs found in *Rhinolophus* bats

We detected SARSr-CoVs in Chinese water myotis (*My. laniger*), Thomas’s horseshoe bat (*Rh. cf. thomasi*) and Thai horseshoe bats (*Rh. siamensis*) from Kunming city (Table S10). While SARSr-CoVs have been identified in bats of the genus *Rhinolophus* before [38–41], to our knowledge, it is the first identification of SARSr-CoVs in *My. laniger*, despite previous analysis of the group [42–44]. This is also important as *Myotis* is more capable of dispersal, including over open environments than *Rhinolophus*, thus it has a disproportionate ability to enable the spread of pathogens across greater distances, or new regions than other known hosts [45, 46]. The SARSr-CoVs in this work were found to be phylogenetically close to Rf4092 (GenBank: KY417145.1) suggesting their evolution from a common ancestor (Figure 4). Rf4092, which was characterized from *Rh. ferrumequinum* in 2012 in Kunming city of Yunnan Province, shares a more recent common ancestor with the SARSr-CoVs identified in this study than with SARS-CoV. It does, however possess considerable sequence identity to SARS-CoV (similarity in the variable region is ORF8 (97.5%), ORF3b (95.6%) and S1 (63.3%) and S2 (95.3%)) [47]. Previous studies suggested host-switching of CoVs between bats and showed ACE2 of mouse-eared bats to enable SARS-CoV and SARS-CoV-2 pseudovirus entry. There was, however, no evidence of β-CoV host switching between mouse-eared bats and horseshoe bats [48, 49]. Due to the close phylogenetic proximity of Rf4092 to the SARSr-CoVs identified here, it is possible that host switching of β-CoV between *Myotis* and *Rhinolophus* bats does occur.

### Some individuals contained more than one bat-related CoV

We observed the coexistence of α-CoV in individual bats (Table S8), which has also been observed previously [21, 50]. It is, however, unclear whether and how co-infection impacts specifically on spillover, evolution and recombination of CoVs. Coexistence of different CoV species in individuals may enable recombination, which when coupled with random mutation enables the emergence of novel lineages of CoVs, further contributing to the diversity of CoVs circulating in bats. Beside the coexistence of CoVs, different virus genera and families were detected in the SARSr-CoVs positive NGS libraries (Figure S2). Although it is difficult to determine their specific relationships with CoVs, most of the cooccurring genera were of *Parvoviridae* origin.

### CoV prevalence was not associated with bat ectoparasite presence

Parasitism in mammals is associated with other physical symptoms of disease. While they are understudied, bat ectoparasites are often shown to contain viruses which they may transmit either actively or passively by mechanical transfer [51], and may also be an indicator of poorer health [52, 53]). However, we did not find any definite trends between bats with ectoparasites and CoV prevalence (Figure S3 and Table S12) [54]. The types and incidence of ectoparasites within different species may be influential and should be considered in the future, both in terms of understanding their roles in bat health status in general, and their roles in transmitting viruses between potential hosts [55].

### Further investigation is necessary to understand SARSr-CoVs in Yunnan Province as well as the ecology of coronaviruses in this region

Phylogenetic analysis of the 11 SARSr-CoV identified here based on the targeted RdRp showed them to be more closely related to SARS-CoV than to SARS-CoV-2 (Figure 4-A). The pathogenicity of these SARSr-CoVs were not determined. We however showed from analysis of NGS data that contigs in 10/11 SARSr-CoV samples covered from 35.5% to 99.4% of the best-hit reference. Isolation of these viruses would enable assessment of their capacity to infect humans, or other mammals through ACE2 receptor binding assays [27, 56]. It will also be important to derive the full-length genome of the SARSr-CoV identified in the *My. laniger*, as it will shed light on the evolution, mutation and recombination of CoVs in nature, which is key to understanding its incidence in the new host.

Dynamics of pathogenic infection in hosts may modulate spillover risk and surveillance of other specimen types e.g., serum, bat guano or urine may offer a supplementary method to assess the risk of spillover. A seasonal trend exists in coronavirus prevalence among isolated bat species [57, 58], and there is extreme variation (0%-80%) in the shedding of bat-borne CoVs through various phases of the breeding season [59, 60], thus surveillance of CoV prevalence in bats with seasonal changes and longitudinally through successive life cycle stages should be prioritized in future research, especially with different patterns of roosting, ecophysiological stressors (such as breeding) and hibernation; all of which impact on susceptibility to contract various pathogens. In addition to a longer period of sampling through successive seasons, a larger sample size over a greater geographical area than Yunnan province would be beneficial. Larger sample sizes can facilitate the assessment of spillover risk within bat communities concerning diversity [61]. Multiple ecological, biological and social factors simultaneously influence coronavirus spillover risk in nature [34, 62–64]. Therefore future modelling of coronavirus prevalence and spillover risk should incorporate these internal and external factors on the background of a much larger sample size.

In conclusion, this research takes us a step closer to understanding the dynamics of CoVs in Asian bats. Not only do we find a broader range of bat hosts of SARSr-CoVs, with the identification in *Myotis laniger*, but we found that bats have the ability to host more than a single CoV concurrently, providing the potential for recombination and the generation of further novel CoVs. Furthermore, we found that within and between species and sites as well as sex, there was a considerable ability for bats to host different CoVs, highlighting the need for more work to understand the mechanisms behind CoV transmission and evolution between bats, and how this varies across space and time. Although our samples come only from Yunnan province and cover a limited period, we find a diversity of CoVs and the prevalence at this specific location and time, reinforcing the need for further sustained work across the region. This work also reiterates the potential for the emergence of novel CoVs from bats, due to the interface between bats and animals with the potential to encounter humans.

## Materials and Methods

### Ethical approval and sample collection

The ethical approval for bat sample collection in Yunnan Province was provided by the Ethics Committee of Life Sciences, Kunming Institute of Zoology (KIZ), Chinese Academy of Sciences (CAS). The ethical approval number is SMKX-20200210-01.

Rectal swabs were collected from bats in 10 different caves in Kunming City (sites: A-H), Yuxi City (site: J) and Xishuangbanna Prefecture (site: I) in Yunnan Province, China, from September to November 2020. Further details of samples collection are provided in Supplementary Method 1.

### Nucleic acid extraction from bat rectal swabs

The samples were retrieved from −80°C, thawed at 4°C, vortexed for 3-5 min and centrifuged at 17 000× *g*, 4℃ for 3 min. The supernatant was transferred into a 0.45 µM filter microtube (Corning, NY, USA), then centrifuged at 15 000× *g*, 4℃ for 1 min. The flow-through was used for nucleic acid purification. Nucleic acids were extracted using the GeneJet Viral DNA and RNA Purification Kit (Thermo Fisher Scientific, MA, USA) and host depletion done using enzyme cocktail digestion [65, 66]. Further details on extraction are in Supplementary Method 2.

### cDNA synthesis and pan-CoV semi-nested PCR detection

cDNA synthesis was performed with TAKARA PrimerScript^TM^ 1st Strand cDNA Synthesis Kit (TAKARA, Dalian, China), following manufacturer instructions. The pan-CoV semi-nested PCR detection consisted of two sets of PCR, the pan-CoV outer PCR and the pan-CoV inner PCR. The primers (synthesized by Tsingke, Beijing, China) are described in Supplementary Method 3. Further details of cDNA synthesis and PCR are in Supplementary Method 4. After PCR, the results were observed by DNA agarose gel electrophoresis and checked by Sanger sequencing, for which details are given in Supplementary Method 5.

### BLAST and analysis of coronavirus prevalence and diversity

The SnapGene software was used to analyze the Sanger sequencing results, by aligning the forward and reverse sequences and electropherogram analysis. After removing the primer sequences, BLASTn was performed on NCBI for all sequences successively. CoVs-positive individuals were identified based on query of targeted RdRp sequences using BLASTn and the best hit virus species were retrieved for subsequent analyses. TaxonKit was used to obtain lineage information for virus species. CoV positive and negative individuals were sorted by different bat species and genera as well as sampling sites in Microsoft Excel (2016). The overlap between the number of bats with ectoparasites and the number of bats with CoVs or without CoVs were analyzed and displayed in Venn diagrams by online tool *Bioinformatics* (https://www.bioinformatics.com.cn/).

### Statistical analysis

Statistical analysis was performed in SPSS (version 24.0). The count for each category as a frequency variate was weighted before descriptive statistics. Confidence intervals (95% CI) were calculated. Crosstabs analysis was used for Chi-square tests. Goodman-Kruskal’s lambda and Cramer’s V coefficient were measured for association analysis. Bonferroni method was chosen to adjust p-values. Different letters (a, b and c) indicate statistically significant differences below the p-value = 0.05 threshold.

### Library preparation for Next generation sequencing (NGS)

Illumina sequencing: Whole transcriptome amplification was performed before library preparation. The WTA2 whole transcriptome amplification (WTA) kit (Sigma-Aldrich, MA, USA) was used for reverse transcription and cDNA synthesis based on the protocol supplied. NGS libraries were prepared by NEBNext^®^ Ultra^™^ II DNA Library Prep Kit for Illumina (NEB, MA, USA) following manufacturer instructions and sequencing was conducted by NovaSeq 6000 (PE150) (Novogene, Shanghai, China).

MGI sequencing: The amplification product of pan-CoV semi-nested PCR was purified and concentrated by the Monarch PCR & DNA Cleanup Kit (NEB), according to the protocol supplied. The libraries were prepared using the MGIEasy FS DNA Library Prep kit (BGI, Shenzhen, China) following manufacturer instructions. The library was sent to BGI for library cyclization and sequencing, and the MGISEQ-2000>PE150 was requested.

### Bioinformatics analysis on NGS data

All NGS data were deposited to the open access database NMDC, China National Microbiology Data Center (https://nmdc.cn/en) (Table S13). Illumina libraries were prepared from samples found positives in the SARSr-CoVs detection. All reads were trimmed by Trimgalore and filtered based on the SILVA database. Contigs were assembled from the reads using SPAdes. Local blastn (Blast+ suite) was utilized to obtain matches in a local blast database consisting of all 2865 CoV sequences initially employed for Pan-CoV PCR primers design using the assembled contigs as queries (contigs in each library). Contig mapping was performed in Geneious R9.

NGS data from Illumina was also processed for the detection of vertebrate-infecting viruses [67]: After read trimming by Trimmomatic, clean reads were aligned to a local database of genomic and transcriptional sequences of bats with Bowtie2. The reads that did not align against the bat sequences were aligned to the human genome. After removing bat and human sequences, de novo assembly was performed with the remaining reads using the MEGAHIT software. The contigs were clustered with CD-HIT-EST software requiring 99% identity and 100% coverage for the shortest sequence. The representative sequences of each cluster were aligned with BLASTN against the Reference Viral Database (RVDB) version 22. The best hit was selected according to e-value, identity, and coverage, in this order of importance. The taxonomic classification of the contigs was assigned according to the taxonomy of the best hit. In parallel, the MEGAN blast2lca tool determined the taxonomy of the contigs according to the lowest common ancestor (LCA) criteria. This information was combined by manual curation and the taxonomies for vertebrate-infecting viruses obtained. The contigs were re-grouped according to the target sequence of the best hit and realigned with the LASTZ tool. From this realignment, the coverage was estimated with Samtools. The contigs were also aligned with Diamond software (The pipeline was implemented by Dr. Alix Armero). The heatmap for vertebrate-host virus was drawn by Graphpad.

MGI sequencing data was utilized for verification of CoV co-infection. Analysis of the amplicon obtained after performing a nested PCR of targeted RdRp region of the CoV genomes consisted of two steps: 1) to estimate the diversity of existing CoVs sequences by the algorithm Emu [68] in conjunction with a fresh homemade genomic CoVs database (accession numbers for the sequences that comprise the genomic CoVs database are provided in an accompanying .txt file as supporting Information). The local CoVs database consists of CoV genome sequences downloaded from GenBank [69] and Virus Pathogen Resource (ViPR) [70] and duplicated sequences were removed; 2) generate the amplification sequence of RdRp regions of CoVs in each sample using an in-house script relying on Minimap2 [71] and alignment produced by Emu in the first step.

### Phylogenetic analysis

The phylogenetic analysis was performed based on the 195 bp sanger sequences obtained from the amplicons from Pan-CoV semi-nested PCR targeting the RdRp sequences of SARSr-CoVs. The same 195 bp RdRp region of reference sequences was also used in the phylogenetic tree. α-CoVs were designated as the outgroups of branches. All of these DNA sequences were aligned by ClustalW algorithm [72]. Models with the lowest BIC scores (Bayesian Information Criterion) [73] are considered to best describe the substitution pattern. The alignment and the best model selection was conducted in MEGA11 [74]. The phylogenetic tree was performed by MEGA11 and QIAGEN CLC Genomics Workbench 22.0.2. The node support was estimated from 1000 bootstrap replicates [75].

## Acknowledgments

We thank Canping Huang for participating in field sampling, and design of pan-CoV primers. We thank Alix Armero for preliminary analysis of the NGS data. We also thank Shengjie Shao, Jialei Chen, Jihao Li for supporting sample collection. We gratefully acknowledge all researchers for sharing their genome data on GISAID. This project was supported by the Ministry of Science and Technology of China (grant no. 2021YFC0863400, 2022YFE0114700), the Alliance of International Scientific Organizations (grant no. ANSO-CR-SP-2020-02), the Shanghai Municipal Science and Technology Major Project (grant no. 2019SHZDZX02), G4 funding from Institut Pasteur, Fondation Merieux and Chinese Academy of Sciences to G.W., and the International Affairs Department of the Institut Pasteur of Paris. A.T. is supported by the ANSO Scholarship for Young Talents. Y.K. is supported by the CAS-TWAS Fellowship for International Doctoral Students.

## Author contributions

**Conceptualization and funding acquisition:** GW and NB. **Supervision:** GW, NB and ACH. **Samples collection:** XC, YC, SX, YK, AT, RL and ACH. **Sample processing:** RL, JY, VO, LM and RX. **Extraction, screening, data analysis and figures drawing:** RL. **Library preparation:** RL, MF, RX and KMB. **Bioinformatics analysis:** AT, NB and SDD. **Writing (original draft):** RL. **Writing (review, editing and revision):** RL, AT, ACH, NB and GW.

