## supplementary material for "Differential prevalence and risk factors for infection with coronaviruses in bats collected during 2020 in Yunnan Province, China"

### Supplementary methods

#### Method S1. Samples collection, bat identification and cave locations

Samples were collected from bats in 10 different locations: site A (sampling date: 19/09/2020), site B (23/09/2020), site C (01/10/2020), site D (30/09/2020), site E (02/10/2020), site F (28/09/2020), site G (29/09/2020), site H (24/09/2020), site I (26/11/2020) and site J (21/09/2020). Bats were collected using 4 bank harp-traps, as well as mistnets, though the number and distributions of these traps and nets varied depending on site specific factors. Nets were set to be ready 30 min before sunset, and left open until activity levels dropped to zero new bats in a 1 h period (typically between 9 pm to midnight). Harp traps were initially checked 30 min after sunset and then checked approximately every 30 min, whereas mistnets were monitored almost constantly. Captured bats were removed and stored in a cloth bag for processing. Species were identified based on our pre-existing analysis which had been validated through barcoding, as well as detailed comparisons of detailed physiology and acoustics [1]. Measurements were made with digital callipers Mitutoyo Absolute Series-500 (accuracy 0.01 mm) and body mass was measured with Pesola Spring Scale (Pesola®Präzisionswaagen AG) with a precision of 0.3%) and included forearm length, head-body, tibia, tail, ear height, and for rhinolophoid bats noseleaf height and width. In addition, profile photos were taken of all individuals, and tissue samples were removed from the wing membrane using a 3mm biopsy punch and stored in 99.7% ethanol.

Rectal swabs from individual bats were collected and the samples were stored in RNAlater (Thermo Fisher Scientific, MA, USA) and transported to the laboratory in dry ice, then stored at -80°C until further processing. The presence, number and identity of ectoparasites on bats were recorded during sampling in site A, site B, site C, site E, site F and site H. Baidu Coordinate Picker (https://api.map.baidu.com/lbsapi/getpoint/) was used to obtain the specific geographical coordinates for the sampling locations, and verified using satellite imagery data in ArcMap.

**Method S2. Nucleic acid extraction from bat rectal swabs**

The extraction was performed according to the protocol supplied with the following modifications: 2 times the volume (400 μL) of input sample solution was used for obtaining a potentially higher viral load. As such, the lysis buffer and proteinase K used were correspondingly doubled in volume. The nucleic acid was eluted with 50 μL RNase-free water (Invitrogen, MA, USA), and stored at -80°C pending further processing.

**Method S3. Degenerate PCR primer design for coronaviruses detection**

Coronavirus sequences were downloaded from NCBI and analyzed in Geneious R9. After sorting, protein sequences were aligned to find the conserved motif of RdRp as the target domain and 1360 RdRp sequences of coronaviruses were extracted and aligned for primer design. During the primer design, inverse degeneration was performed following the codon list [2]. The sequence up to approximately 20 bp upstream from the target was also selected as a clamp sequence to complete three universal primers for pan-coronavirus (pan-CoV) semi-nested PCR detection (during primer synthesis, the N residues in the primer sequences were replaced with hypoxanthine (I)). The sequences of the pan-CoV outer semi-nested PCR primers (target size 298 bp) were as follows: Pan-CoVs-OF: 5'-TGTTATTGGAACAACTAAATTYTAYGGNGGNTG-3' (position 1900 to 1933 in alignment) and Pan-CoVs-OR: 5'-GGTTGCATCACCACTACTAGTNCCNCCNGGYTT-3' (position 2165 to 2197 in alignment); The pan-CoVs inner semi-nested PCR primer (target size 236 bp) was as follows: Pan-CoVs-IF: 5'-GTTTTGAAAATCCTATTCTTATGGGNTGGGAYTAYCC-3' (position 1962 to 1998 in alignment), whereas the Pan-CoVs-OR was used as the reverse primer. The alignment is provided in an extended txt file. Sanger sequencing was performed with the following primers: CoVs-seq-OR: 5'-GGTTGCATCACCACTACT-3' and CoVs-seq-IF: 5'-TGAAAATCCTATTCTTATG-3'.

**Method S4. cDNA synthesis and pan-CoV semi-nested PCR detection**

Random hexamers (50 μM) were used in this kit as random primers in synthesis. The protocol is as follows: reverse transcription at 30°C for 10 min, then 42°C for 45 min. Enzyme denaturation at 95°C for 5 min, before proceeding to PCR.

The pan-CoV outer PCR mix is as follows: 10 μL TAKARA Ex Taq Version 2.0 plus dye (TAKARA, Dalian, China), 1 μL Pan-CoVs-OF (10 μM) as forward primer, 1 μL Pan-CoVs-OR (10 μM) as reverse primer, 2 μL cDNA, and nuclease-free water was added to make a total reaction volume of 20 μL. The pan-CoV outer PCR protocol is as follows: Initial denaturation at 94°C for 2 min, 5 cycles (denaturation at 94°C for 30 s, annealing at 44°C for 30 s and extension at 72°C for 20 s), 30 cycles (denaturation at 94°C for 30 s, annealing at 54°C for 30 s and extension at 72°C for 20 s), followed by a final extension at 72°C for 2 min and hold at 4°C.

The pan-CoV inner PCR mix is as follows: 10 μL TAKARA Ex Taq Version 2.0 plus dye, 1 μL Pan-CoVs-IF (10 μM) as forward primer, 1 μL Pan-CoVs-OR (10 μM) as reverse primer, 0.8 μL outer PCR products, 1 μL 5% DMSO (Yeasen Biotech, Shanghai, China) and nuclease-free water was added to make a total reaction volume of 20 μL. The pan-CoV inner PCR protocol is as follows: Initial denaturation at 94°C for 2 min, 5 cycles (denaturation at 94°C for 30 s, annealing at 50°C for 30 s and extension at 72°C for 20 s), 30 cycles (denaturation at 94°C for 30 s, annealing at 58°C for 30 s and extension at 72°C for 20 s), followed by a final extension at 72°C for 2 min and hold at 4°C.

**Method S5. DNA agarose gel electrophoresis and Sanger sequencing**

1.5% agarose (Baygene, Shanghai, China) gel was in 1×TAE buffer (Sangon, Shanghai, China) for DNA electrophoresis. A concentration of 0.1% nucleic acid dye Tanon, Shanghai, China) was added to the agarose gel to visualize the DNA. 3 μL of DL5000 Marker (Tanon, Shanghai, China) was used and 6 μL of sample were loaded into each well. The DNA agarose gel electrophoresis (AGE) was run at 110 V for 40 min. After the AGE run was complete, a gel image was taken to identify a 236 bp band at the expected targeted size. When the band of the expected size was present, 3 tubes for PCR were prepared for each positive template, and the AGE was performed again as described above. The target bands were excised under UV light for Sanger sequencing (Tsingke, Beijing, China). Bidirectional sequencing with reverse primer (CoVs-seq-OR) and forward primers (CoVs-seq-IF) were performed to enable nested PCR for all positive templates.

**Method S6. Identity verification of bats with SARSr-CoVs**

Using two pairs of primers (inner and outer) for nested PCR amplification, bat species were identified by molecular barcoding, targeting the conserved gene cytochrome c oxidase subunit I (COI). The primer (Tsingke) sequences [3] are as follows: Outer nested PCR primers: BEGLCOIf: 5'- GGYGCYTGAGCHGGWATAGT-3' and BEGLCOIr: 5'- ARRATDGGRTCYCCYCCTCC-3'. The reaction consists of 10 μL TAKARA Ex Taq Version 2.0 plus dye (TAKARA, Dalian, China), 1 μL BEGLCOIf (10 μM) as forward primer, 1 μL BEGLCOIr (10 μM) as reverse primer, 1.5 μL cDNA and nuclease-free water to a total volume of 20 μL. The reaction conditions are: Initial denaturation at 94°C for 2 min, 35 cycles (denaturation at 94°C for 30 s, annealing at

50°C for 30 s and extension at 72°C for 50 s), final extension at 72°C for 2 min and hold at 4°C. Inner nested PCR primers: SFF_145f: 5’-GTHACHGCYCAYGCHTTYGTAATAAT-3’ and SFF_351r: 5’-CTCCWGCRTGDGCWAGRTTTCC-3’. The reaction consists of 10 μL TAKARA Ex Taq Version 2.0 plus dye, 1 μL SFF_145f (10 μM) as forward primer, 1 μL SFF_351r (10 μM) as reverse primer, 0.5 μL outer PCR products, 1 μL 5% DMSO (Yeasen Biotech, Shanghai, China) and nuclease-free water to a total volume of 20 μL. The reaction conditions are: Initial denaturation at 94°C for 2 min, 35 cycles (denaturation at 94°C for 30 s, annealing at 53°C for 30 s and extension at 72°C for 20 s), final extension at 72°C for 2 min and hold at 4°C.

Human coronavirus 229E (HCoV-229E) and a null template control were utilized as positive and negative controls, respectively, in each batch of nested PCR and agarose gel electrophoresis.

### Supplementary figures

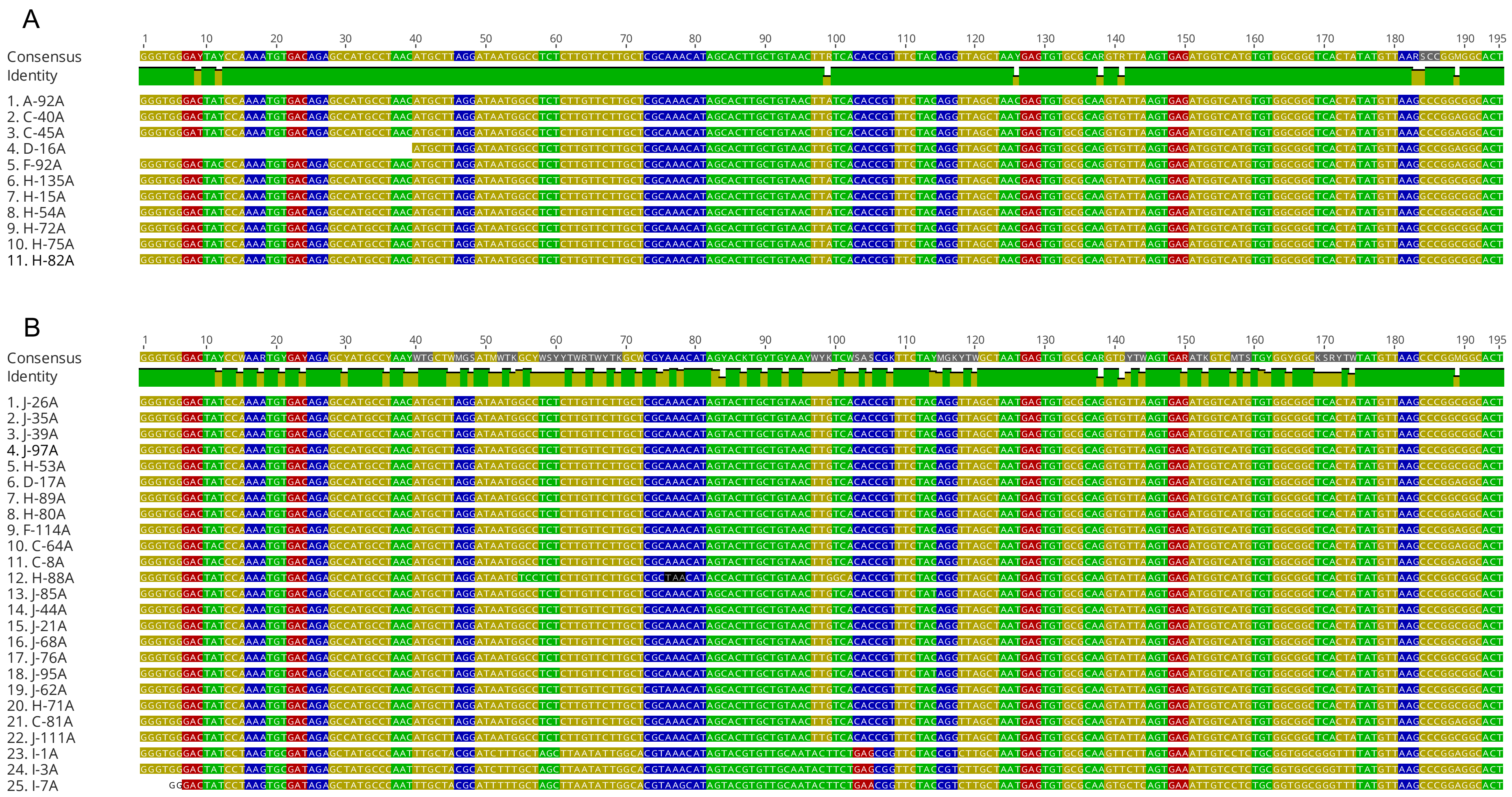

C

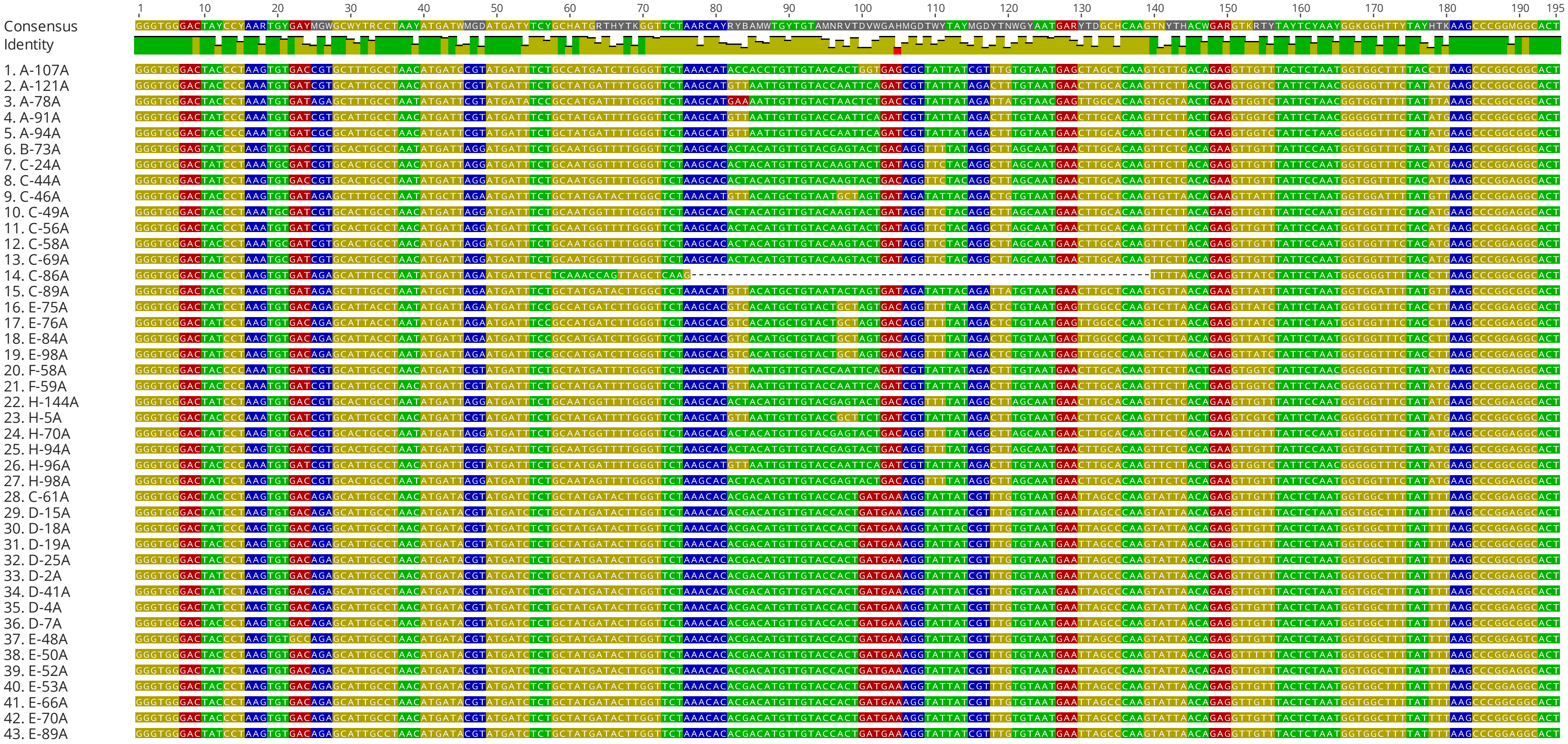
1 10 20 30 40 50 60 70 80 90 100 110 120 130 140 150 160 170 180 190 195
Consensus
Identity
1. A-107A
2. A-121A
3. A-78A
4. A-91A
5. A-94A
6. B-73A
7. C-24A
8. C-44A
9. C-46A
10. C-49A
11. C-56A
12. C-58A
13. C-69A
14. C-86A
15. C-89A
16. E-75A
17. E-76A
18. E-84A
19. E-98A
20. F-58A
21. F-59A
22. H-144A
23. H-5A
24. H-70A
25. H-94A
26. H-96A
27. H-98A
28. C-61A
29. D-15A
30. D-18A
31. D-19A
32. D-25A
33. D-2A
34. D-41A
35. D-4A
36. D-7A
37. E-48A
38. E-50A
39. E-52A
40. E-53A
41. E-66A
42. E-70A
43. E-89A

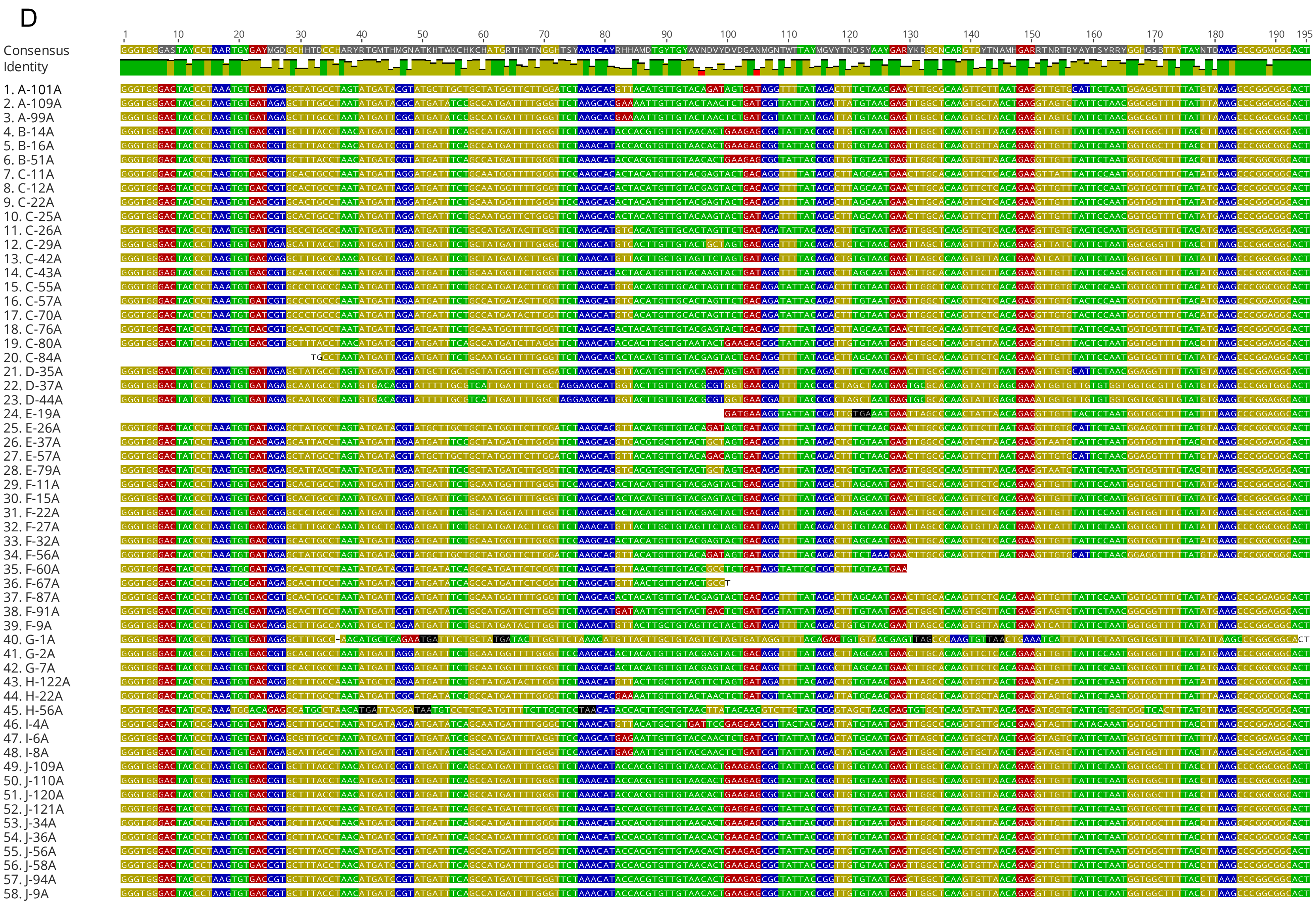
D
Consensus
Identity
1. A-101A
2. A-109A
3. A-99A
4. B-14A
5. B-16A
6. B-51A
7. C-11A
8. C-12A
9. C-22A
10. C-25A
11. C-26A
12. C-29A
13. C-42A
14. C-43A
15. C-55A
16. C-57A
17. C-70A
18. C-76A
19. C-80A
20. C-84A
21. D-35A
22. D-37A
23. D-44A
24. E-19A
25. E-26A
26. E-37A
27. E-57A
28. E-79A
29. F-11A
30. F-15A
31. F-22A
32. F-27A
33. F-32A
34. F-56A
35. F-60A
36. F-67A
37. F-87A
38. F-91A
39. F-9A
40. G-1A
41. G-2A
42. G-7A
43. H-122A
44. H-22A
45. H-56A
46. I-4A
47. I-6A
48. I-8A
49. J-109A
50. J-110A
51. J-120A
52. J-121A
53. J-34A
54. J-36A
55. J-56A
56. J-58A
57. J-94A
58. J-9A

**Figure S1. Alignment of 195bp targeted RdRp sequences of coronaviruses from 137 CoV-positive individuals.** (A) Alignment for 11 SARSr-CoV positives. (B) Alignment for 25 Beta-CoVs positives, excluding the 11 SARSr-CoVs. (C) Alignment for 43 Alpha-CoV positives. (D) Alignment for 58 unclassified CoV positives. All the sequences are available by the accessions in NMDC attached in a txt file 'sequences ID and accessions' and '137 targeted RdRp sequences' in Supporting Information.

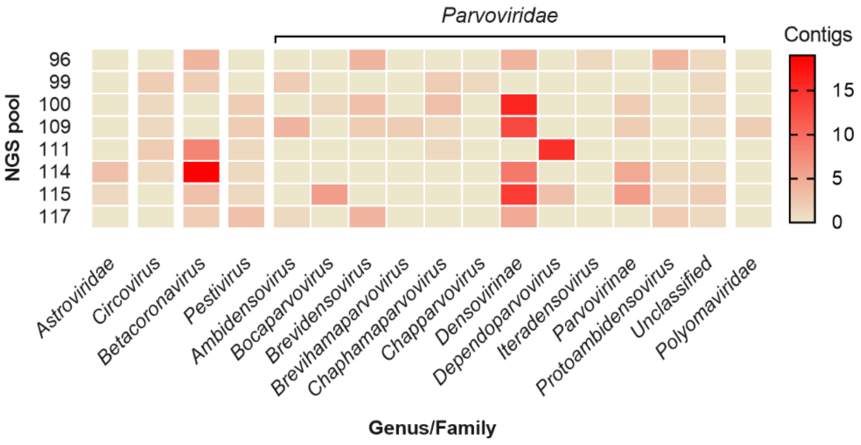


**Figure S2. Heat map of contigs of vertebrate-infecting viruses in the Illumina sequencing libraries including the 11 SARSr-CoV positive individuals.** The virome is shown by genus, if it is unclassified, shown by the family.

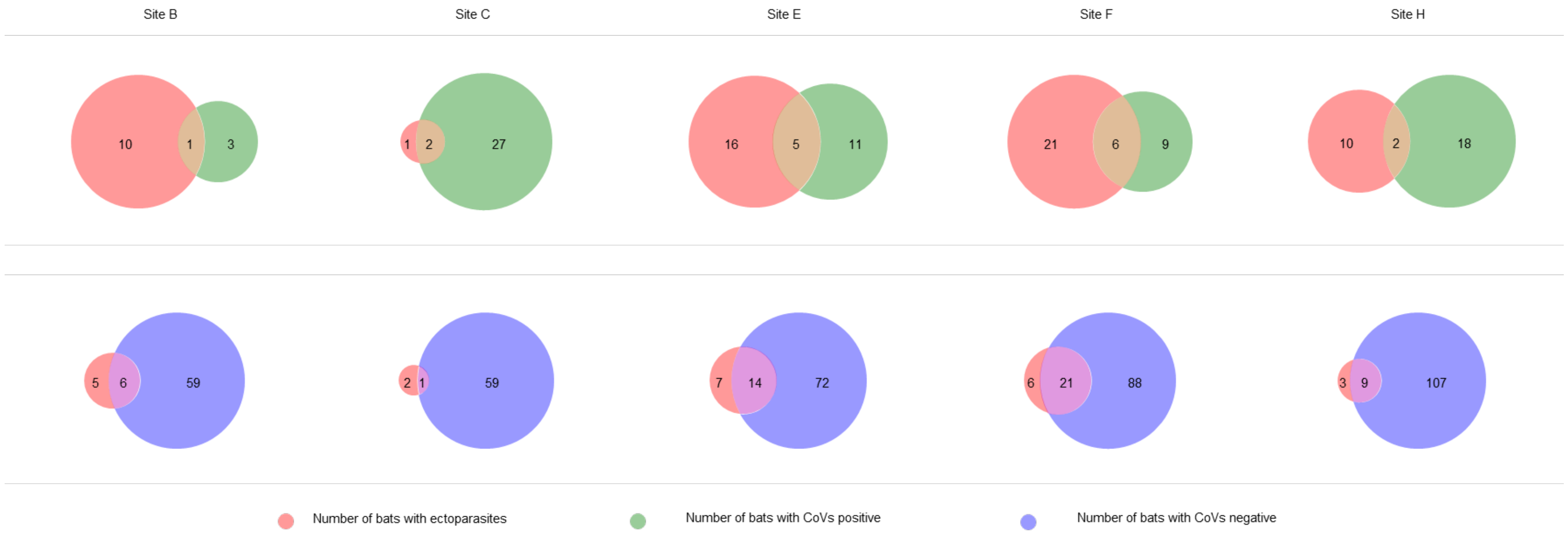


**Figure S3. The Venn diagram of bats with CoVs or without CoVs and bats with ectoparasites in different locations.** The light red colour indicates the number of bats with ectoparasites. The light green colour indicates the number of bats with CoVs positive. The light violet indicates the number of bats with CoVs negative. The light brown indicates the overlap between the number of bats with ectoparasites and bats with CoVs. The pink colour indicates the overlap between the number of bats with ectoparasites and bats without CoVs.

### Supplementary tables

#### Table S1. Coronavirus prevalence in rectal swabs from bats in Yunnan province, China.

| Species \ Sampling sites | No. of CoV-positive samples/total no. of samples; CoV-prevalence of sampling site[a]: | | | | | | | | | | |
|---|---|---|---|---|---|---|---|---|---|---|---|
| | **Kunming** | | | | | | | | **Xishuangbanna** | **Yuxi** | **All locations** |
| | **A** | **B** | **C*** | **D*** | **E** | **F** | **G^** | **H** | **I^** | **J*** | |
| *E. spelaea*^ | 0/0; NA | 0/0; NA | 0/0; NA | 0/0; NA | 0/0; NA | 0/0; NA | 0/0; NA | 0/0; NA | 2/2; 100.0%^ | 0/0; NA | 2/2; 100.0%^ |
| *H. armiger* | 0/0; NA | 0/0; NA | 1/5; 20.0% | 0/0; NA | 0/12; 0.0% | 0/1; 0.0% | 0/0; NA | 1/4; 25.0% | 0/0; NA | 0/5; 0.0% | 2/27; 7.4% |
| *H. pomona* | 0/1; 0.0% | 0/0; NA | 1/1; 100.0%^ | 10/44; 22.7% | 8/48; 16.7% | 0/2; 0.0% | 0/0; NA | 0/0; NA | 0/0; NA | 0/0; NA | 19/96; 19.8% |
| *I. io* | 0/1; 0.0% | 0/0; NA | 0/0; NA | 0/0; NA | 0/0; NA | 0/0; NA | 0/0; NA | 0/0; NA | 0/0; NA | 0/0; NA | 0/1; 0.0% |
| *Me. lyra*^ | 0/0; NA | 0/0; NA | 0/0; NA | 0/0; NA | 0/0; NA | 0/0; NA | 0/0; NA | 0/0; NA | 1/2; 50.0%^ | 0/0; NA | 1/2; 50.0% |
| *Mi. schreibersii** | 6/7; 85.7%^ | 0/0; NA | 0/0; NA | 0/0; NA | 0/0; NA | 5/34; 14.7% | 0/0; NA | 3/6; 50.0%^ | 2/3; 66.7%^ | 0/10; 0.0% | 16/60; 26.7%* |
| *My. fimbriatus* | 0/0; NA | 1/9; 11.1% | 9/20; 45.0%* | 0/0; NA | 0/0; NA | 6/21; 28.6%* | 3/5; 60.0%^ | 5/49; 10.2% | 0/0; NA | 0/0; NA | 24/104; 23.1% |
| *My. laniger* | 0/0; NA | 0/37; 0.0% | 14/46; 30.4%* | 0/0; NA | 0/0; NA | 0/3; 0.0% | 0/1; 0.0% | 0/1; 0.0% | 0/0; NA | 0/4; 0.0% | 14/92; 15.2% |
| *My. pilosus* | 0/0; NA | 0/0; NA | 0/0; NA | 0/0; NA | 6/19; 31.6%* | 0/6; 0.0% | 0/0; NA | 0/1; 0.0% | 0/0; NA | 0/0; NA | 6/26; 23.1% |
| *My. siligorensis*^ | 0/0; NA | 0/1; 0.0% | 0/0; NA | 0/0; NA | 0/0; NA | 0/0; NA | 0/0; NA | 0/0; NA | 0/0; NA | 1/1; 100.0%^ | 1/2; 50.0% |
| *My. sp.* | 0/3; 0.0% | 0/0; NA | 0/0; NA | 0/0; NA | 0/0; NA | 0/0; NA | 0/0; NA | 0/0; NA | 0/0; NA | 0/0; NA | 0/3; 0.0% |
| *Rh. affinis* | 0/22; 0.0% | 0/0; NA | 0/1; 0.0% | 1/2; 50.0%^ | 2/23; 8.7% | 1/29; 3.4% | 0/2; 0.0% | 0/0; NA | 0/0; NA | 0/17; 0.0% | 4/96; 4.2% |
| *Rh. pusillus* | 0/0; NA | 0/0; NA | 0/2; 0.0% | 0/0; NA | 0/0; NA | 0/0; NA | 0/0; NA | 0/1; 0.0% | 0/0; NA | 0/5; 0.0% | 0/8; 0.0% |
| *Rh. rex* | 0/0; NA | 0/0; NA | 0/0; NA | 0/0; NA | 0/0; NA | 0/3; 0.0% | 0/0; NA | 0/0; NA | 0/0; NA | 0/6; 0.0% | 0/9; 0.0% |
| *Rh. siamensis* | 0/0; NA | 0/0; NA | 0/6; 0.0% | 0/0; NA | 0/0; NA | 0/0; NA | 0/0; NA | 1/8; 12.5% | 0/0; NA | 0/11; 0.0% | 1/25; 4.0% |
| *Rh. sinicus** | 3/11; 27.3%^ | 3/22; 13.6% | 4/8; 50.0%^ | 2/2; 100.0%^ | 0/0; NA | 3/25; 12.0% | 0/0; NA | 10/66; 15.2% | 0/1; 0.0% | 21/27; 77.8%* | 46/162; 28.4%* |
| *Ro. amplexicaudatus*^ | 0/0; NA | 0/0; NA | 0/0; NA | 0/0; NA | 0/0; NA | 0/0; NA | 0/0; NA | 0/0; NA | 1/1; 100.0%^ | 0/0; NA | 1/1; 100.0%^ |
| **All species** | 9/45; 20.0% | 4/69; 5.8% | 29/89; 32.6%* | 13/48; 27.1%* | 16/102; 15.7% | 15/124; 12.1% | 3/8; 37.5%^ | 20/136; 14.7% | 6/9; 66.7%^ | 22/86; 25.6%* | 137/716; 19.1% |

[a]NA, No bats collected. Notable samples are indicated by * if there are over 15 bats with an incidence of over 25% and ^ notes a high percentage, but the small sample-size means this may not be representative.

**Table S2. Differential prevalence of coronaviruses as distributed between bat sex.**

| Bat Species | Alpha/Prevalence | | | Beta/Prevalence | | | Unclassified/Prevalence | | | Count/ CoV prevalence | | | Count (All) | | |
|---|---|---|---|---|---|---|---|---|---|---|---|---|---|---|---|
| | F | M | Sum | F | M | Sum | F | M | Sum | F | M | Sum | F | M | Sum |
| *E. spelaea* | 0 | 0 | 0 | 1/100% | 1/100% | 2/100% | 0 | 0 | 0 | 1/100% | 1/100% | 2/100% | 1 | 1 | 2 |
| *H armiger* | 0 | 0 | 0 | 2/10.5% | 0 | 2/7.4% | 0 | 0 | 0 | 2/10.5% | 0 | 2/7.4% | 19 | 8 | 27 |
| *H. pomona* | 8/14.3% | 8/21.1% | 16/17.0% | 0 | 0 | 0 | 1/1.8% | 2/5.3% | 3/3.2% | 9/16.1% | 10/26.3% | 19/20.2% | 56 | 38 | 94 |
| *Me. lyra* | NA | 0 | 0 | NA | 0 | 0 | NA | 1/50.0% | 1/50.0% | NA | 1/50% | 1/50% | NA | 2 | 2 |
| *Mi. schreibersii* | 3/9.7% | 1/5.0% | 4/7.0% | 0 | 0 | 0 | 3/9.7% | 3/15.0% | 6/11.8% | 6/19.4% | 4/20.0% | 10/19.6% | 31 | 20 | 51 |
| *My. fimbriatus* | 1/1.9% | 4/8.2% | 5/4.9% | 1/1.9% | 0 | 1/1.0% | 9/16.7% | 9/18.4% | 18/17.5% | 11/20.4% | 13/26.5% | 24/23.3% | 54 | 49 | 103 |
| *My. laniger* | 5/10.0% | 3/7.1% | 8/8.7% | 0 | 1/2.4% | 1/1.1% | 1/2.0% | 4/9.5% | 5/5.4% | 6/12.0% | 8/19.0% | 14/15.2% | 50 | 42 | 92 |
| *My. pilosus* | 0 | 4/18.2% | 4/15.4% | 0 | 0 | 0 | 0 | 2/9.1% | 2/7.7% | 0 | 6/27.3% | 6/23.1% | 4 | 22 | 26 |
| *My. siligorensis* | 0 | 0 | 0 | 0 | 1/100% | 1/50.0% | 0 | 0 | 0 | 0 | 1/100% | 1/50% | 1 | 1 | 2 |
| *Rh. affinis* | 0 | 0 | 0 | 0 | 12/46.2% | 12/16.4% | 1/2.1% | 3/11.5% | 4/5.5% | 1/2.1% | 3/11.5% | 4/5.5% | 47 | 26 | 73 |
| *Rh. siamensis* | 0 | 0 | 0 | 1/6.7% | 0 | 1/4.3% | 0 | 0 | 0 | 1/6.7% | 0 | 1/4.3% | 15 | 8 | 23 |
| *Rh. sinicus* | 1/1.1% | 0 | 1/0.7% | 14/16.1% | 0 | 14/9.3% | 10/11.5% | 6/9.5% | 16/10.7% | 25/28.7% | 18/28.6% | 43/28.7% | 87 | 63 | 150 |
| *Ro. amplexicaudatus* | NA | 0 | 0 | NA | 1/100% | 1/100% | NA | 0 | 0 | NA | 1/100% | 1/100% | NA | 1 | 1 |
| Total | 18 | 20 | 38 | 19 | 16 | 35 | 25 | 30 | 55 | 62/16.5% | 66/23.0% | 128/19.3% | 376 | 287 | 663 |

NA, No bats collected.

**Table S3. Differential prevalence of coronavirus in different bat species and sampling sites.**

| Species \ Sampling sites | No. of α-CoV positive samples/no. of β-CoV positive samples/no. of unclassified *Coronavirinae* positive samples*: | | | | | | | | | | |
|---|---|---|---|---|---|---|---|---|---|---|---|
| | **Kunming** | | | | | | | | **Xishuangbanna** | **Yuxi** | **All locations** |
| | **A** | **B** | **C** | **D** | **E** | **F** | **G** | **H** | **I** | **J** | |
| *E. spelaea* | — | — | — | — | — | — | — | — | 0/2/0 | — | 0/2/0 |
| *H. armiger* | — | — | 0/1/0 | — | — | — | — | 0/1/0 | — | — | 0/2/0 |
| *H. pomona* | — | — | 1/0/0 | 8/0/2 | 7/0/1 | — | — | — | — | — | 16/0/3 |
| *Me. lyra* | — | — | — | — | — | — | — | — | 0/0/1 | — | 0/0/1 |
| *Mi. schreibersii* | 4/0/2 | — | — | — | — | 2/0/3 | — | 2/0/1 | 0/0/2 | — | 8/0/8 |
| *My. fimbriatus* | — | 1/0/0 | 1/0/8 | — | — | 0/0/6 | 0/0/3 | 3/1/1 | — | — | 5/1/18 |
| *My. laniger* | — | — | 8/1/5 | — | — | — | — | — | — | — | 8/1/5 |
| *My. pilosus* | — | — | — | — | 4/0/2 | — | — | — | — | — | 4/0/2 |
| *My. siligorensis* | — | — | — | — | — | — | — | — | — | 0/1/0 | 0/1/0 |
| *Rh. affinis* | — | — | — | 0/0/1 | 0/0/2 | 0/0/1 | — | — | — | — | 0/0/4 |
| *Rh. siamensis* | — | — | — | — | — | — | — | 0/1/0 | — | — | 0/1/0 |
| *Rh. sinicus* | 1/1/1 | 0/0/3 | 0/3/1 | 0/2/0 | — | 0/2/1 | — | 1/8/1 | — | 0/11/10 | 2/27/17 |
| *Ro. amplexicaudatus* | — | — | — | — | — | — | — | — | 0/1/0 | — | 0/1/0 |
| **All species** | 5/1/3 | 1/0/3 | 10/5/14 | 8/2/3 | 11/0/5 | 2/2/11 | 0/0/3 | 6/11/3 | 0/3/3 | 0/12/10 | 43/36/58 |

*—, No prevalence of coronavirus.

**Table S4. Sex of sampled bats as distributed across species.**

| Bat Species | Bat sex | | | Total |
|---|---|---|---|---|
| | Female | Male | Unidentified sex | |
| *E. spelaea* | 1 | 1 | 0 | 2 |
| *H armiger* | 19 | 8 | 0 | 27 |
| *H. pomona* | 56 | 38 | 2 | 96 |
| *I. io* | NA | NA | 1 | 1 |
| *Me. lyra* | NA | 2 | 0 | 2 |
| *Mi. schreibersii* | 31 | 20 | 9 | 60 |
| *My. fimbriatus* | 54 | 49 | 1 | 104 |
| *My. laniger* | 50 | 42 | 0 | 92 |
| *My. pilosus* | 4 | 22 | 0 | 26 |
| *My. siligorensis* | 1 | 1 | 0 | 2 |
| *My. sp.* | NA | NA | 3 | 3 |
| *Rh. affinis* | 47 | 26 | 23 | 96 |
| *Rh. pusillus* | 5 | 3 | 0 | 8 |
| *Rh. rex* | 6 | 3 | 0 | 9 |
| *Rh. siamensis* | 15 | 8 | 2 | 25 |
| *Rh. sinicus* | 87 | 63 | 12 | 162 |
| *Ro. amplexicaudatus* | NA | 1 | 0 | 1 |
| Total | 376 | 287 | 53 | 716 |

NA, No bats collected.

**Table S5. BLASTn results for CoV-positive samples.**

| No. | Sample ID | Bat species | Sex | Query length (bp) | Identity % | Accession (Genbank) | Genus | Subgenus | Virus species/strain |
|---|---|---|---|---|---|---|---|---|---|
| 1 | D-19A | *H. pomona* | F | 195 | 94.9 | MN611523.1 | *Alphacoronavirus* | *Decacovirus* | *H. pomona bat coronavirus HKU10-related* |
| 2 | D-25A | *H. pomona* | F | 197 | 96.72 | MN611523.1 | *Alphacoronavirus* | *Decacovirus* | *H. pomona bat coronavirus HKU10-related* |
| 3 | D-7A | *H. pomona* | F | 195 | 94.9 | MN611523.1 | *Alphacoronavirus* | *Decacovirus* | *H. pomona bat coronavirus HKU10-related* |
| 4 | E-66A | *H. pomona* | F | 195 | 94.9 | MN611523.1 | *Alphacoronavirus* | *Decacovirus* | *H. pomona bat coronavirus HKU10-related* |
| 5 | E-70A | *H. pomona* | F | 195 | 94.9 | MN611523.1 | *Alphacoronavirus* | *Decacovirus* | *H. pomona bat coronavirus HKU10-related* |
| 6 | C-61A | *H. pomona* | M | 195 | 94.9 | MN611523.1 | *Alphacoronavirus* | *Decacovirus* | *H. pomona bat coronavirus HKU10-related* |
| 7 | E-48A | *H. pomona* | M | 195 | 96.17 | MN611523.1 | *Alphacoronavirus* | *Decacovirus* | *H. pomona bat coronavirus HKU10-related* |

| No. | Sample ID | Bat species | Sex | Query length (bp) | Identity % | Accession (Genbank) | Genus | Subgenus | Virus species/strain |
|---|---|---|---|---|---|---|---|---|---|
| 8 | E-50A | *H. pomona* | F | 195 | 94.39 | MN611523.1 | *Alphacoronavirus* | *Decacovirus* | *H. pomona bat coronavirus HKU10-related* |
| 9 | E-52A | *H. pomona* | M | 195 | 94.39 | MN611523.1 | *Alphacoronavirus* | *Decacovirus* | *H. pomona bat coronavirus HKU10-related* |
| 10 | E-53A | *H. pomona* | M | 195 | 94.9 | MN611523.1 | *Alphacoronavirus* | *Decacovirus* | *H. pomona bat coronavirus HKU10-related* |
| 11 | E-89A | *H. pomona* | M | 196 | 94.42 | MN611523.1 | *Alphacoronavirus* | *Decacovirus* | *H. pomona bat coronavirus HKU10-related* |
| 12 | D-18A | *H. pomona* | F | 195 | 93.88 | MN611523.1 | *Alphacoronavirus* | *Decacovirus* | *H. pomona bat coronavirus HKU10-related* |
| 13 | D-41A | *H. pomona* | M | 195 | 94.39 | MN611523.1 | *Alphacoronavirus* | *Decacovirus* | *H. pomona bat coronavirus HKU10-related* |
| 14 | D-4A | *H. pomona* | F | 195 | 94.39 | MN611523.1 | *Alphacoronavirus* | *Decacovirus* | *H. pomona bat coronavirus HKU10-related* |
| 15 | D-15A | *H. pomona* | M | 195 | 94.39 | MN611523.1 | *Alphacoronavirus* | *Decacovirus* | *H. pomona bat coronavirus HKU10-related* |
| 16 | D-2A | *H. pomona* | M | 195 | 94.39 | MN611523.1 | *Alphacoronavirus* | *Decacovirus* | *H. pomona bat coronavirus HKU10-related* |

| No. | Sample ID | Bat species | Sex | Query length (bp) | Identity % | Accession (Genbank) | Genus | Subgenus | Virus species/strain |
|---|---|---|---|---|---|---|---|---|---|
| 17 | A-121A | *Mi. schreibersii* | NA | 195 | 97.45 | KJ473800.1 | *Alphacoronavirus* | *Minunacovirus* | *Miniopterus bat coronavirus HKU8* |
| 18 | A-91A | *Mi. schreibersii* | NA | 195 | 96.94 | KJ473800.1 | *Alphacoronavirus* | *Minunacovirus* | *Miniopterus bat coronavirus HKU8* |
| 19 | A-94A | *Mi. schreibersii* | NA | 195 | 96.94 | KJ473800.1 | *Alphacoronavirus* | *Minunacovirus* | *Miniopterus bat coronavirus HKU8* |
| 20 | H-5A | *Mi. schreibersii* | F | 195 | 95.41 | KJ473800.1 | *Alphacoronavirus* | *Minunacovirus* | *Miniopterus bat coronavirus HKU8* |
| 21 | H-96A | *Mi. schreibersii* | F | 195 | 97.45 | KJ473800.1 | *Alphacoronavirus* | *Minunacovirus* | *Miniopterus bat coronavirus HKU8* |
| 22 | F-58A | *Mi. schreibersii* | M | 195 | 97.45 | KJ473800.1 | *Alphacoronavirus* | *Minunacovirus* | *Miniopterus bat coronavirus HKU8* |
| 23 | F-59A | *Mi. schreibersii* | F | 195 | 97.45 | KJ473800.1 | *Alphacoronavirus* | *Minunacovirus* | *Miniopterus bat coronavirus HKU8* |
| 24 | C-89A | *My. laniger* | M | 195 | 97.73 | KU973689.1 | *Alphacoronavirus* | *unclassified Alphacoronavirus* | *Alphacoronavirus sp.* |
| 25 | B-73A | *My. fimbriatus* | M | 195 | 94.62 | MZ081397.1 | *Alphacoronavirus* | *unclassified Alphacoronavirus* | *Alphacoronavirus sp.* |
| 26 | C-24A | *My. laniger* | F | 195 | 96.24 | MZ081397.1 | *Alphacoronavirus* | *unclassified Alphacoronavirus* | *Alphacoronavirus sp.* |
| 27 | C-44A | *My. laniger* | F | 195 | 96.24 | MZ081397.1 | *Alphacoronavirus* | *unclassified Alphacoronavirus* | *Alphacoronavirus sp.* |
| 28 | C-46A | *My. laniger* | F | 195 | 98.3 | KU973689.1 | *Alphacoronavirus* | *unclassified Alphacoronavirus* | *Alphacoronavirus sp.* |
| 29 | C-49A | *My. laniger* | F | 195 | 95.7 | MZ081397.1 | *Alphacoronavirus* | *unclassified Alphacoronavirus* | *Alphacoronavirus sp.* |

| No. | Sample ID | Bat species | Sex | Query length (bp) | Identity % | Accession (Genbank) | Genus | Subgenus | Virus species/strain |
|---|---|---|---|---|---|---|---|---|---|
| 30 | C-56A | *My. laniger* | M | 195 | 96.24 | MZ081397.1 | *Alphacoronavirus* | *unclassified Alphacoronavirus* | *Alphacoronavirus sp.* |
| 31 | C-58A | *My. laniger* | F | 195 | 95.7 | MZ081397.1 | *Alphacoronavirus* | *unclassified Alphacoronavirus* | *Alphacoronavirus sp.* |
| 32 | C-69A | *My. laniger* | M | 195 | 96.24 | MZ081397.1 | *Alphacoronavirus* | *unclassified Alphacoronavirus* | *Alphacoronavirus sp.* |
| 33 | H-70A | *Rh. sinicus* | F | 195 | 94.62 | MZ081397.1 | *Alphacoronavirus* | *unclassified Alphacoronavirus* | *Alphacoronavirus sp.* |
| 34 | H-144A | *My. fimbriatus* | M | 195 | 94.62 | MZ081397.1 | *Alphacoronavirus* | *unclassified Alphacoronavirus* | *Alphacoronavirus sp.* |
| 35 | H-94A | *My. fimbriatus* | M | 195 | 94.62 | MZ081397.1 | *Alphacoronavirus* | *unclassified Alphacoronavirus* | *Alphacoronavirus sp.* |
| 36 | H-98A | *My. fimbriatus* | M | 196 | 94.12 | MZ081397.1 | *Alphacoronavirus* | *unclassified Alphacoronavirus* | *Alphacoronavirus sp.* |
| 37 | E-75A | *My. pilosus* | M | 195 | 98.36 | DQ249224.1 | *Alphacoronavirus* | *unclassified Alphacoronavirus* | *Bat coronavirus HKU6* |
| 38 | E-76A | *My. pilosus* | M | 195 | 98.36 | DQ249224.1 | *Alphacoronavirus* | *unclassified Alphacoronavirus* | *Bat coronavirus HKU6* |
| 39 | E-84A | *My. pilosus* | M | 195 | 98.36 | DQ249224.1 | *Alphacoronavirus* | *unclassified Alphacoronavirus* | *Bat coronavirus HKU6* |
| 40 | E-98A | *My. pilosus* | M | 195 | 98.36 | DQ249224.1 | *Alphacoronavirus* | *unclassified Alphacoronavirus* | *Bat coronavirus HKU6* |
| 41 | A-78A | *Mi. schreibersii* | NA | 196 | 98.37 | KJ473804.1 | *Alphacoronavirus* | *unclassified Alphacoronavirus* | *BtMf-AlphaCoV/HeN2013-a* |
| 42 | A-107A | *Rh. sinicus* | NA | 195 | 96.43 | MK211373.1 | *Alphacoronavirus* | *unclassified Alphacoronavirus* | *Coronavirus BtRs-AlphaCoV/YN2018* |

| No. | Sample ID | Bat species | Sex | Query length (bp) | Identity % | Accession (Genbank) | Genus | Subgenus | Virus species/strain |
|---|---|---|---|---|---|---|---|---|---|
| 43 | C-86A | *My. fimbriatus* | F | 132 | 89.74 | KP895494.1 | *Alphacoronavirus* | *unclassified Alphacoronavirus* | *Myotis Bat Alphacoronavirus strain YNXY_46C* |
| 44 | I-1A | *Ro. amplexicaudatus* | M | 195 | 96.94 | KU762338.1 | *Betacoronavirus* | *Nobecovirus* | *Rousettus bat coronavirus GCCDC1* |
| 45 | I-3A | *E. spelaea* | F | 195 | 96.94 | KU762338.1 | *Betacoronavirus* | *Nobecovirus* | *Rousettus bat coronavirus GCCDC1* |
| 46 | I-7A | *E. spelaea* | M | 191 | 96.34 | KU762337.1 | *Betacoronavirus* | *Nobecovirus* | *Rousettus bat coronavirus GCCDC1* |
| 47 | H-15A | *Rh. cf. thomasi* | F | 195 | 98.91 | KY417149.1 | *Betacoronavirus* | *Sarbecovirus* | *Severe acute respiratory syndrome-related coronavirus* |
| 48 | A-92A | *Rh. sinicus* | NA | 195 | 98.91 | KY417149.1 | *Betacoronavirus* | *Sarbecovirus* | *Severe acute respiratory syndrome-related coronavirus* |
| 49 | H-54A | *Rh. sinicus* | F | 195 | 98.91 | KY417149.1 | *Betacoronavirus* | *Sarbecovirus* | *Severe acute respiratory syndrome-related coronavirus* |
| 50 | H-72A | *Rh. sinicus* | M | 195 | 98.91 | KY417149.1 | *Betacoronavirus* | *Sarbecovirus* | *Severe acute respiratory* |

| No. | Sample ID | Bat species | Sex | Query length (bp) | Identity % | Accession (Genbank) | Genus | Subgenus | Virus species/strain |
|---|---|---|---|---|---|---|---|---|---|
| | | | | | | | | | *syndrome-related coronavirus* |
| 51 | H-75A | *Rh. siamensis* | F | 195 | 98.91 | KY417149.1 | *Betacoronavirus* | *Sarbecovirus* | *Severe acute respiratory syndrome-related coronavirus* |
| 52 | F-92A | *Rh. sinicus* | F | 195 | 98.91 | FJ588686.1 | *Betacoronavirus* | *Sarbecovirus* | *Severe acute respiratory syndrome-related coronavirus* |
| 53 | H-135A | *Rh. sinicus* | F | 195 | 98.91 | KY417149.1 | *Betacoronavirus* | *Sarbecovirus* | *Severe acute respiratory syndrome-related coronavirus* |
| 54 | H-82A | *Rh. sinicus* | F | 195 | 98.91 | KY417149.1 | *Betacoronavirus* | *Sarbecovirus* | *Severe acute respiratory syndrome-related coronavirus* |
| 55 | C-40A | *Rh. sinicus* | M | 195 | 99.45 | FJ588686.1 | *Betacoronavirus* | *Sarbecovirus* | *Severe acute respiratory syndrome-related coronavirus* |
| 56 | C-45A | *My. laniger* | M | 195 | 99.46 | KY417145.1 | *Betacoronavirus* | *Sarbecovirus* | *Severe acute respiratory syndrome-related coronavirus* |

| No. | Sample ID | Bat species | Sex | Query length (bp) | Identity % | Accession (Genbank) | Genus | Subgenus | Virus species/strain |
|---|---|---|---|---|---|---|---|---|---|
| 57 | D-16A | *Rh. sinicus* | M | 156 | 100 | KY417145.1 | *Betacoronavirus* | *Sarbecovirus* | *Severe acute respiratory syndrome-related coronavirus* |
| 58 | J-26A | *Rh. sinicus* | F | 195 | 98.91 | OK017857.1 | *Betacoronavirus* | *Sarbecovirus* | *unclassified Sarbecovirus* |
| 59 | J-35A | *Rh. sinicus* | F | 195 | 98.91 | OK017857.1 | *Betacoronavirus* | *Sarbecovirus* | *unclassified Sarbecovirus* |
| 60 | J-39A | *Rh. sinicus* | F | 195 | 98.91 | OK017857.1 | *Betacoronavirus* | *Sarbecovirus* | *unclassified Sarbecovirus* |
| 61 | J-97A | *Rh. sinicus* | F | 195 | 98.91 | OK017857.1 | *Betacoronavirus* | *Sarbecovirus* | *unclassified Sarbecovirus* |
| 62 | J-85A | *Rh. sinicus* | F | 195 | 98.91 | MK211377.1 | *Betacoronavirus* | *Sarbecovirus* | *Coronavirus BtRs-BetaCoV/YN2018C* |
| 63 | J-44A | *Rh. sinicus* | M | 195 | 98.91 | MK211378.1 | *Betacoronavirus* | *Sarbecovirus* | *Coronavirus BtRs-BetaCoV/YN2018D* |
| 64 | J-111A | *Rh. sinicus* | M | 195 | 98.91 | MK211378.1 | *Betacoronavirus* | *Sarbecovirus* | *Coronavirus BtRs-BetaCoV/YN2018D* |
| 65 | J-21A | *Rh. sinicus* | M | 195 | 98.91 | MK211378.1 | *Betacoronavirus* | *Sarbecovirus* | *Coronavirus BtRs-BetaCoV/YN2018D* |
| 66 | J-68A | *Rh. sinicus* | M | 195 | 98.91 | MK211378.1 | *Betacoronavirus* | *Sarbecovirus* | *Coronavirus BtRs-BetaCoV/YN2018D* |
| 67 | J-76A | *Rh. sinicus* | M | 195 | 98.91 | MK211378.1 | *Betacoronavirus* | *Sarbecovirus* | *Coronavirus BtRs-BetaCoV/YN2018D* |
| 68 | J-95A | *Rh. sinicus* | M | 195 | 98.91 | MK211377.1 | *Betacoronavirus* | *Sarbecovirus* | *Coronavirus BtRs-BetaCoV/YN2018C* |

| No. | Sample ID | Bat species | Sex | Query length (bp) | Identity % | Accession (Genbank) | Genus | Subgenus | Virus species/strain |
|---|---|---|---|---|---|---|---|---|---|
| 69 | J-62A | *My. siligorensis* | M | 195 | 98.36 | MK211378.1 | *Betacoronavirus* | *Sarbecovirus* | *Coronavirus BtRs-BetaCoV/YN2018D* |
| 70 | H-53A | *Rh. sinicus* | F | 195 | 98.91 | OK017857.1 | *Betacoronavirus* | *Sarbecovirus* | *unclassified Sarbecovirus* |
| 71 | H-71A | *H. armiger* | F | 195 | 98.91 | MK211378.1 | *Betacoronavirus* | *Sarbecovirus* | *Coronavirus BtRs-BetaCoV/YN2018D* |
| 72 | F-114A | *Rh. sinicus* | M | 195 | 98.91 | OK017857.1 | *Betacoronavirus* | *Sarbecovirus* | *unclassified Sarbecovirus* |
| 73 | H-88A | *Rh. sinicus* | M | 196 | 94.57 | MK211378.1 | *Betacoronavirus* | *Sarbecovirus* | *Coronavirus BtRs-BetaCoV/YN2018D* |
| 74 | H-89A | *Rh. sinicus* | F | 195 | 98.91 | OK017857.1 | *Betacoronavirus* | *Sarbecovirus* | *unclassified Sarbecovirus* |
| 75 | C-64A | *Rh. sinicus* | F | 195 | 98.36 | OK017857.1 | *Betacoronavirus* | *Sarbecovirus* | *unclassified Sarbecovirus* |
| 76 | C-8A | *Rh. sinicus* | F | 195 | 98.36 | OK017857.1 | *Betacoronavirus* | *Sarbecovirus* | *unclassified Sarbecovirus* |
| 77 | D-17A | *Rh. sinicus* | M | 195 | 98.91 | OK017857.1 | *Betacoronavirus* | *Sarbecovirus* | *unclassified Sarbecovirus* |
| 78 | H-80A | *Rh. sinicus* | F | 196 | 98.37 | OK017857.1 | *Betacoronavirus* | *Sarbecovirus* | *unclassified Sarbecovirus* |
| 79 | C-81A | *H. armiger* | F | 195 | 98.91 | MK211378.1 | *Betacoronavirus* | *Sarbecovirus* | *Coronavirus BtRs-BetaCoV/YN2018D* |
| 80 | A-109A | *Mi. schreibersii* | NA | 195 | 96.17 | KF294280.1 | *unclassified Coronavirinae* | *Bat coronavirus* | *NA* |
| 81 | A-99A | *Mi. schreibersii* | NA | 195 | 96.72 | KF294280.1 | *unclassified Coronavirinae* | *Bat coronavirus* | *NA* |

| No. | Sample ID | Bat species | Sex | Query length (bp) | Identity % | Accession (Genbank) | Genus | Subgenus | Virus species/strain |
|---|---|---|---|---|---|---|---|---|---|
| 82 | H-22A | *Mi. schreibersii* | F | 195 | 98.91 | KF294280.1 | *unclassified Coronavirinae* | *Bat coronavirus* | *NA* |
| 83 | J-56A | *Rh. sinicus* | F | 195 | 95.08 | MG916902.1 | *unclassified Coronavirinae* | *Bat coronavirus* | *NA* |
| 84 | J-9A | *Rh. sinicus* | F | 195 | 95.08 | MG916902.1 | *unclassified Coronavirinae* | *Bat coronavirus* | *NA* |
| 85 | J-110A | *Rh. sinicus* | F | 195 | 95.63 | MG916902.1 | *unclassified Coronavirinae* | *Bat coronavirus* | *NA* |
| 86 | J-120A | *Rh. sinicus* | F | 195 | 95.08 | MG916902.1 | *unclassified Coronavirinae* | *Bat coronavirus* | *NA* |
| 87 | J-94A | *Rh. sinicus* | F | 195 | 95.16 | MG916902.1 | *unclassified Coronavirinae* | *Bat coronavirus* | *NA* |
| 88 | J-121A | *Rh. sinicus* | M | 195 | 94.54 | MG916902.1 | *unclassified Coronavirinae* | *Bat coronavirus* | *NA* |
| 89 | J-34A | *Rh. sinicus* | M | 195 | 95.08 | MG916902.1 | *unclassified Coronavirinae* | *Bat coronavirus* | *NA* |
| 90 | J-58A | *Rh. sinicus* | M | 195 | 95.08 | MG916902.1 | *unclassified Coronavirinae* | *Bat coronavirus* | *NA* |
| 91 | J-109A | *Rh. sinicus* | M | 195 | 95.08 | MG916902.1 | *unclassified Coronavirinae* | *Bat coronavirus* | *NA* |
| 92 | J-36A | *Rh. sinicus* | M | 195 | 95.08 | MG916902.1 | *unclassified Coronavirinae* | *Bat coronavirus* | *NA* |
| 93 | B-14A | *Rh. sinicus* | F | 195 | 95.08 | MG916902.1 | *unclassified Coronavirinae* | *Bat coronavirus* | *NA* |
| 94 | B-16A | *Rh. sinicus* | F | 195 | 95.08 | MG916902.1 | *unclassified Coronavirinae* | *Bat coronavirus* | *NA* |

| No. | Sample ID | Bat species | Sex | Query length (bp) | Identity % | Accession (Genbank) | Genus | Subgenus | Virus species/strain |
|---|---|---|---|---|---|---|---|---|---|
| 95 | B-51A | *Rh. sinicus* | M | 195 | 95.08 | MG916902.1 | *unclassified Coronavirinae* | *Bat coronavirus* | *NA* |
| 96 | H-56A | *Rh. sinicus* | F | 195 | 80.32 | KY780391.1 | *unclassified Coronavirinae* | *Bat coronavirus* | *Bat coronavirus Rfer4019* |
| 97 | F-60A | *Mi. schreibersii* | F | 129 | 96.12 | KF294277.1 | *unclassified Coronavirinae* | *Bat coronavirus* | *NA* |
| 98 | F-67A | *Mi. schreibersii* | F | 100 | 97.03 | KF294277.1 | *unclassified Coronavirinae* | *Bat coronavirus* | *NA* |
| 99 | F-91A | *Mi. schreibersii* | M | 196 | 89.94 | KF294273.1 | *unclassified Coronavirinae* | *Bat coronavirus* | *NA* |
| 100 | C-80A | *Rh. sinicus* | F | 195 | 93.44 | MG916902.1 | *unclassified Coronavirinae* | *Bat coronavirus* | *NA* |
| 101 | D-37A | *H. pomona* | F | 195 | 96.41 | MZ969008.1 | *unclassified Coronavirinae* | *Bat coronavirus* | *NA* |
| 102 | D-44A | *H. pomona* | M | 195 | 95.9 | MZ969008.1 | *unclassified Coronavirinae* | *Bat coronavirus* | *NA* |
| 103 | E-37A | *My. pilosus* | M | 195 | 100 | DQ648825.1 | *unclassified Coronavirinae* | *Bat coronavirus China 2005* | *Bat coronavirus (BtCoV/A604/2005)* |
| 104 | E-79A | *My. pilosus* | M | 195 | 99.42 | DQ648825.1 | *unclassified Coronavirinae* | *Bat coronavirus China 2005* | *Bat coronavirus (BtCoV/A604/2005)* |
| 105 | I-6A | *Mi. schreibersii* | M | 195 | 97.27 | KU343189.1 | *unclassified Coronavirinae* | *Bat coronavirus MsBtCoV/3710* | *NA* |
| 106 | I-8A | *Mi. schreibersii* | M | 195 | 98.36 | KU343189.1 | *unclassified Coronavirinae* | *Bat coronavirus MsBtCoV/3710* | *NA* |
| 107 | E-57A | *Rh. affinis* | F | 195 | 97.45 | KU343198.1 | *unclassified Coronavirinae* | *Bat coronavirus RaBtCoV/3750* | *NA* |

| No. | Sample ID | Bat species | Sex | Query length (bp) | Identity % | Accession (Genbank) | Genus | Subgenus | Virus species/strain |
|---|---|---|---|---|---|---|---|---|---|
| 108 | F-56A | *Rh. affinis* | M | 195 | 96.43 | KU343198.1 | *unclassified Coronavirinae* | *Bat coronavirus RaBtCoV/3750* | *NA* |
| 109 | D-35A | *Rh. affinis* | M | 195 | 97.96 | KU343198.1 | *unclassified Coronavirinae* | *Bat coronavirus RaBtCoV/3750* | *NA* |
| 110 | E-26A | *Rh. affinis* | M | 195 | 95.41 | KU343199.1 | *unclassified Coronavirinae* | *Bat coronavirus RaBtCoV/4307-2* | *NA* |
| 111 | A-101A | *Rh. sinicus* | NA | 195 | 95.41 | KU343199.1 | *unclassified Coronavirinae* | *Bat coronavirus RaBtCoV/4307-2* | *NA* |
| 112 | C-55A | *My. laniger* | M | 195 | 79.08 | JQ731782.1 | *unclassified Coronavirinae* | *Coronavirus BtCoV/KP816/Phy_dis/PAN/2011* | *NA* |
| 113 | C-26A | *My. laniger* | M | 195 | 79.08 | JQ731782.1 | *unclassified Coronavirinae* | *Coronavirus BtCoV/KP816/Phy_dis/PAN/2011* | *NA* |
| 114 | C-57A | *My. laniger* | M | 195 | 79.08 | JQ731782.1 | *unclassified Coronavirinae* | *Coronavirus BtCoV/KP816/Phy_dis/PAN/2011* | *NA* |
| 115 | C-70A | *My. laniger* | M | 195 | 79.08 | JQ731782.1 | *unclassified Coronavirinae* | *Coronavirus BtCoV/KP816/Phy_dis/PAN/2011* | *NA* |
| 116 | I-4A | *Me. lyra* | M | 195 | 84.02 | GU065427.1 | *unclassified Coronavirinae* | *Kenya bat coronavirus BtKY83* | *NA* |
| 117 | C-29A | *My. fimbriatus* | F | 195 | 96.11 | KF569978.1 | *unclassified Coronavirinae* | *Myotis daubentonii coronavirus* | *NA* |

| No. | Sample ID | Bat species | Sex | Query length (bp) | Identity % | Accession (Genbank) | Genus | Subgenus | Virus species/strain |
|---|---|---|---|---|---|---|---|---|---|
| 118 | C-12A | *My. fimbriatus* | F | 195 | 97.06 | KF569991.1 | *unclassified Coronavirinae* | *Myotis davidii coronavirus* | *NA* |
| 119 | C-22A | *My. fimbriatus* | F | 195 | 96.47 | KF569991.1 | *unclassified Coronavirinae* | *Myotis davidii coronavirus* | *NA* |
| 120 | C-42A | *My. fimbriatus* | F | 195 | 99.4 | KF569993.1 | *unclassified Coronavirinae* | *Myotis davidii coronavirus* | *NA* |
| 121 | C-11A | *My. fimbriatus* | M | 195 | 95.88 | KF569991.1 | *unclassified Coronavirinae* | *Myotis davidii coronavirus* | *NA* |
| 122 | C-25A | *My. fimbriatus* | M | 195 | 98.82 | KF569991.1 | *unclassified Coronavirinae* | *Myotis davidii coronavirus* | *NA* |
| 123 | C-43A | *My. fimbriatus* | M | 195 | 98.24 | KF569991.1 | *unclassified Coronavirinae* | *Myotis davidii coronavirus* | *NA* |
| 124 | C-76A | *My. fimbriatus* | M | 195 | 97.06 | KF569991.1 | *unclassified Coronavirinae* | *Myotis davidii coronavirus* | *NA* |
| 125 | G-1A | *My. fimbriatus* | F | 194 | 99.4 | KF569993.1 | *unclassified Coronavirinae* | *Myotis davidii coronavirus* | *NA* |
| 126 | G-7A | *My. fimbriatus* | F | 195 | 97.06 | KF569991.1 | *unclassified Coronavirinae* | *Myotis davidii coronavirus* | *NA* |
| 127 | G-2A | *My. fimbriatus* | M | 195 | 96.47 | KF569991.1 | *unclassified Coronavirinae* | *Myotis davidii coronavirus* | *NA* |
| 128 | F-22A | *My. fimbriatus* | M | 196 | 92.98 | KF569991.1 | *unclassified Coronavirinae* | *Myotis davidii coronavirus* | *NA* |
| 129 | F-27A | *My. fimbriatus* | M | 195 | 97.6 | KF569993.1 | *unclassified Coronavirinae* | *Myotis davidii coronavirus* | *NA* |
| 130 | F-9A | *My. fimbriatus* | M | 195 | 97.01 | KF569993.1 | *unclassified Coronavirinae* | *Myotis davidii coronavirus* | *NA* |

| No. | Sample ID | Bat species | Sex | Query length (bp) | Identity % | Accession (Genbank) | Genus | Subgenus | Virus species/strain |
|---|---|---|---|---|---|---|---|---|---|
| 131 | H-122A | *My. fimbriatus* | M | 195 | 97.6 | KF569993.1 | *unclassified Coronavirinae* | *Myotis davidii coronavirus* | *NA* |
| 132 | F-11A | *My. fimbriatus* | F | 195 | 96.47 | KF569991.1 | *unclassified Coronavirinae* | *Myotis davidii coronavirus* | *NA* |
| 133 | F-15A | *My. fimbriatus* | F | 195 | 97.06 | KF569991.1 | *unclassified Coronavirinae* | *Myotis davidii coronavirus* | *NA* |
| 134 | F-32A | *My. fimbriatus* | F | 195 | 96.47 | KF569991.1 | *unclassified Coronavirinae* | *Myotis davidii coronavirus* | *NA* |
| 135 | F-87A | *Rh. sinicus* | F | 195 | 97.06 | KF569991.1 | *unclassified Coronavirinae* | *Myotis davidii coronavirus* | *NA* |
| 136 | C-84A | *My. laniger* | F | 164 | 97.4 | KF569991.1 | *unclassified Coronavirinae* | *Myotis davidii coronavirus* | *NA* |
| 137 | E-19A | *H. pomona* | M | 96 | 89.58 | KF569979.1 | *unclassified Coronavirinae* | *Rhinolophus ferrumequinum coronavirus* | *NA* |

Sample ID indicates the information of the sampling site and the bat number and the type of samples (A in the label means rectal swab). Bat sex: F (Famle) and M (Male).

**Table S6. Crosstabs analysis between coronavirus prevalence with different factors and the correlation coefficient.**

| CROSSTABS | Sampling site | Bat genus | Bat species | | Bat sex |
|---|---|---|---|---|---|
| NA | | | Rhinolophus | Myotis | |
| Chi-Square Test[a] | | | | | |
| Pearson $\chi^2$ | 29.791*** | 3.473 | 22.648*** | 2.094 | 4.424* |
| Nonparametric Correlation Coefficient | | | | | |
| Cramer's V | 0.206*** | 0.070 | 0.296*** | 0.097 | 0.082* |
| Goodman-Kruskal tau | 0.043*** | 0.005 | 0.088* | 0.009 | 0.007* |
| N of Valid Cases | 699 | 710 | 258 | 222 | 663 |

*, significance at p-values below the 0.05 threshold. ***, significance at p-values below the 0.001 threshold. a, 0 cells (0.0%) have expected count less than 5 in Chi-Square test.

**Table S7. Differential prevalence of α-, β-, unclassified-CoVacross multiple bat genera.**

| Genera | Alpha | | | Beta | | | Unclassfied | | | Count (Positive) | | |
|---|---|---|---|---|---|---|---|---|---|---|---|---|
| | F | M | Sum | F | M | Sum | F | M | Sum | F | M | Sum |
| *Eonycteris* | NA | NA | 0 | 1 | 1 | 2 | NA | NA | 0 | 1 | 1 | 2 |
| *Hipposideros* | 8 | 8 | 16 | 2 | NA | 2 | 1 | 2 | 3 | 11 | 10 | 21 |
| *Megaderma* | NA | NA | 0 | NA | NA | 0 | NA | 1 | 1 | NA | 1 | 1 |
| *Miniopterus* | 3 | 1 | 4 | NA | NA | 0 | 3 | 3 | 6 | 6 | 4 | 10 |
| *Myotis* | 6 | 11 | 17 | 1 | 2 | 3 | 10 | 15 | 25 | 17 | 28 | 45 |
| *Rhinolophus* | 1 | NA | 1 | 15 | 12 | 27 | 11 | 9 | 20 | 27 | 21 | 48 |
| *Rousettus* | NA | NA | 0 | NA | 1 | 1 | NA | NA | 0 | NA | 1 | 1 |
| **Total** | 18 | 20 | 38 | 19 | 16 | 35 | 25 | 30 | 55 | 62 | 66 | 128 |
| **Genera** | **Alpha-CoV prevalence** | | | **Beta-CoV prevalence** | | | **Unclassified-CoV prevalence** | | | **CoV prevalence** | | |
| | F | M | Sum | F | M | Sum | F | M | Sum | F | M | Sum |
| *Eonycteris* | NA | NA | 0.0% | 100.0 % | 100.0 % | 100.0 % | NA | NA | 0.0% | 100.0 % | 100.0 % | 100.0 % |
| *Hipposideros* | 10.7% | 17.4% | 13.2% | 2.7% | NA | 1.7% | 1.3% | 4.3% | 2.5% | 14.7% | 21.7% | 17.4% |
| *Megaderma* | NA | NA | 0.0% | NA | NA | 0.0% | NA | 50.0% | 50.0% | NA | 50.0% | 50.0% |
| *Miniopterus* | 9.7% | 5.0% | 7.8% | NA | NA | 0.0% | 9.7% | 15.0% | 11.8% | 19.4% | 20.0% | 19.6% |
| *Myotis* | 5.5% | 9.6% | 7.6% | 0.9% | 1.8% | 1.3% | 9.2% | 13.2% | 11.2% | 15.6% | 24.6% | 20.2% |
| *Rhinolophus* | 0.6% | NA | 0.4% | 9.4% | 11.7% | 10.3% | 6.9% | 8.7% | 7.6% | 16.9% | 20.4% | 18.3% |
| *Rousettus* | NA | NA | 0.0% | NA | 100.0 % | 100.0 % | NA | NA | 0.0% | NA | 100.0 % | 100.0 % |
| **Total** | 4.8% | 7.0% | 5.7% | 5.1% | 5.6% | 5.3% | 6.6% | 10.5% | 8.3% | 16.5% | 23.0% | 19.3% |

**Table S8. Differential prevalence of CoV co-existing in individual bat samples.**

| No. | Sample label | DSI % | BLAST TaxID | Genus | Virus species |
|---|---|---|---|---|---|
| 1 | C-29A-1 | 82.6 | 1906673 | *Alphacoronavirus* | *Alphacoronavirus sp.* |
| | C-29A-2 | 13.9 | 1244203 | *Alphacoronavirus* | *Bat coronavirus HKU10* |
| | C-29A-3 | 1.8 | 1508220 | *unclassified Coronavirinae* | *Bat coronavirus* |
| 2 | C-42A-1 | 88.2 | 1487702 | *unclassified Coronavirinae* | *Myotis davidii coronavirus* |
| | C-42A-2 | 7.7 | 1487702 | *unclassified Coronavirinae* | *Myotis davidii coronavirus* |
| | C-42A-3 | 3.3 | 1487701 | *unclassified Coronavirinae* | *Myotis daubentonii coronavirus* |
| 3 | C-86A-1 | 84.2 | 1906673 | *Alphacoronavirus* | *Alphacoronavirus sp.* |
| | C-86A-2 | 10.3 | 393045 | *Alphacoronavirus* | *Bat coronavirus HKU6* |
| | C-86A-3 | 2.2 | 1487701 | *unclassified Coronavirinae* | *Myotis daubentonii coronavirus* |
| | C-86A-4 | 1.7 | 1487701 | *unclassified Coronavirinae* | *Myotis daubentonii coronavirus* |

DSI: Diversity of sequences in individual bats.

**Table S9. Crosstabs analysis between differential prevalence of CoV with different factors and the correlation coefficient.**

| CROSSTABS | Sampling site[a] | Bat genus[b] | Bat species[c] | Bat sex[b] |
|---|---|---|---|---|
| Chi-Square Test | | | | |
| Pearson $\chi^2$ | 46.564*** | 56.203*** | 81.907*** | 0.693 |
| Nonparametric Correlation Coefficient | | | | |
| Cramer's V | 0.450*** | 0.490*** | 0.572*** | 0.074 |
| Goodman-Kruskal tau | 0.197*** | 0.223*** | 0.303 | 0.003 |
| N of Valid Cases | 115 | 117 | 125 | 128 |

*, significance at p-values below the 0.05 threshold. ***, significance at p-values below the 0.001 threshold. a, 5 cells (27.8%), b, 0 cells (0.0%), c, 7 cells (38.9%) have expected count less than 5 in Chi-Square test.

**Table S10. Host verification and virus species of 11 SARSr-CoV positive samples.**

| No. | Sample ID[a] | Query length (bp) | Query cover | Percent identity | Accession (Genbank) | Bat species (COI gene) | Subgenus | Virus species |
|---|---|---|---|---|---|---|---|---|
| 1 | A-92A | 195 | 93% | 98.91 | KY417149.1 | *Rh. cf. thomasi* | *Sarbecovirus* | *Bat SARS-like coronavirus Rs4255* |
| 2 | C-40A | 195 | 93% | 99.45 | FJ588686.1 | *Rh. cf. thomasi* | *Sarbecovirus* | *SARS coronavirus Rs_672/2006* |
| 3 | C-45A | 195 | 94% | 99.46 | KY417145.1 | *My. laniger* | *Sarbecovirus* | *Bat SARS-like coronavirus Rf4092* |
| 4 | D-16A | 156 | 91% | 100 | KY417145.1 | *Rh. cf. thomasi* | *Sarbecovirus* | *Bat SARS-like coronavirus Rf4092* |
| 5 | F-92A | 195 | 93% | 98.91 | FJ588686.1 | *Rh. cf. thomasi* | *Sarbecovirus* | *SARS coronavirus Rs_672/2006* |
| 6 | H-15A | 195 | 93% | 96.34 | KY417149.1 | *Rh. cf. thomasi* | *Sarbecovirus* | *Bat SARS-like coronavirus Rs4255* |
| 7 | H-54A | 195 | 93% | 98.91 | KY417149.1 | *Rh. cf. thomasi* | *Sarbecovirus* | *Bat SARS-like coronavirus Rs4255* |
| 8 | H-72A | 195 | 93% | 98.91 | KY417149.1 | *Rh. cf. thomasi* | *Sarbecovirus* | *Bat SARS-like coronavirus Rs4255* |
| 9 | H-75A | 195 | 93% | 98.91 | KY417149.1 | *Rh. siamensis* | *Sarbecovirus* | *Bat SARS-like coronavirus Rs4255* |
| 10 | H-82A | 195 | 93% | 98.91 | KY417149.1 | *Rh. cf. thomasi* | *Sarbecovirus* | *Bat SARS-like coronavirus Rs4255* |
| 11 | H-135A | 195 | 93% | 98.91 | KY417149.1 | *Rh. cf. thomasi* | *Sarbecovirus* | *Bat SARS-like coronavirus Rs4255* |

a. Sample ID indicates the information of the sampling site and the bat number and the type of samples (A in the label means rectal swab).

**Table S11. Statistics for contigs mapping to whole genome and different genes of SARSr-CoVs.**

| Gene | Whole | | ORF1a | | ORF1b | | S | | 3a | | E | |
|---|---|---|---|---|---|---|---|---|---|---|---|---|
| Size* | 29743 bp | | 13149 bp | | 7887 bp | | 3726 bp | | 825 bp | | 231 bp | |
| Mapping | Coverage | Identity | Coverage | Identity | Coverage | Identity | Coverage | Identity | Coverage | Identity | Coverage | Identity |
| Pool 41 | 18369 | 93.9 | 6543 | 94.2 | 4752 | 93.2 | 3152 | 93.2 | 741 | 96.0 | NA | NA |
| Pool 96 | 20241 | 91.0 | 8773 | 91.2 | 5464 | 91.5 | 2447 | 90.3 | 758 | 94.5 | 168 | 33.1 |
| Pool 100 | 208 | 87.0 | NA | NA | NA | NA | NA | NA | NA | NA | NA | NA |
| Pool 111 | 23539 | 89.6 | 8938 | 91.5 | 6135 | 93.9 | 3726^ | 72.7 | 784 | 86.6 | 231^ | 98.3 |
| Pool 114 | 29567^ | 91.7 | 12973 | 97.8 | 7887^ | 96.3 | 3726^ | 76.7 | 825^ | 89.9 | 231^ | 98.7 |
| Pool 115 | 24095 | 92.6 | 10398 | 94.6 | 6297 | 93.5 | 2832 | 90.0 | 825^ | 87.7 | 231^ | 98.7 |
| Pool 117 | 10565 | 92.0 | 3839 | 92.2 | 1776 | 86.7 | 1573 | 94.5 | 444 | 85.4 | NA | NA |
| Size# | 29059 bp | | 12570 bp | | 7887 bp | | 3726 bp | | 825 bp | | 231 bp | |
| Pool 99 | 2189 | 90.6 | 530 | 91.5 | 952 | 90.4 | NA | NA | NA | NA | NA | NA |
| Pool 109 | 2216 | 86.1 | 777 | 92.3 | 295 | 88.3 | NA | NA | NA | NA | NA | NA |
| Gene | M | | 6 | | 7a | | 7b | | 8 | | N | |
| Size* | 693 bp | | 192 bp | | 369 bp | | 135 bp | | 366 bp | | 1269 bp | |
| Mapping | Coverage | Identity | Coverage | Identity | Coverage | Identity | Coverage | Identity | Coverage | Identity | Coverage | Identity |
| Pool 41 | 557 | 95.3 | 192^ | 97.6 | 208 | 90.4 | 135^ | 98.5 | 366^ | 89.2 | 1269^ | 95.4 |
| Pool 96 | 693^ | 92.8 | 192^ | 98.4 | 123 | 84.6 | NA | NA | 200 | 90.0 | 1149 | 95.0 |
| Pool 100 | NA | NA | NA | NA | NA | NA | NA | NA | NA | NA | 208 | 87.0 |
| Pool 111 | 693^ | 95.0 | 192^ | 99.5 | 369^ | 90.3 | 135^ | 99.3 | 366^ | 97.8 | 1269^ | 97.9 |
| Pool 114 | 693^ | 98.1 | 192^ | 99.5 | 369^ | 98.5 | 135^ | 98.5 | 366^ | 98.9 | 1269^ | 97.9 |
| Pool 115 | 690 | 79.2 | 192^ | 38.1 | 369^ | 97.6 | 135^ | 98.5 | 366^ | 98.1 | 1269^ | 98.2 |
| Pool 117 | 559 | 95.3 | 192^ | 97.4 | 25 | 32.0 | 123 | 87.8 | 366^ | 97.8 | 1265^ | 93.7 |
| Size# | 693 bp | | 192 bp | | 369 bp | | 135 bp | | 366 bp | | 1269 bp | |
| Pool 99 | NA | NA | NA | NA | NA | NA | NA | NA | NA | NA | 707 | 90.3 |
| Pool 109 | NA | NA | NA | NA | NA | NA | NA | NA | 226 | 39.0 | 904 | 91.6 |

*, size of the reference sequence (GenBank: KY417149.1); #, size of the reference sequence (GenBank: FJ588686.1); ^, the coverage for the size of references is more than 99%.

**Table S12. The information of the CoV-positive bats with ectoparasite.**

| No. | Bat ID[a] | Bat species | CoVs positive |
|---|---|---|---|
| 1 | B-14 | *Rh. sinicus* | *unclassified Coronavirinae* |
| 2 | C-22 | *My. fimbriatus* | *unclassified Coronavirinae* |
| 3 | C-55 | *My. laniger* | *unclassified Coronavirinae* |
| 4 | E-37 | *My. pilosus* | *unclassified Coronavirinae* |
| 5 | E-75 | *My. pilosus* | *Alphacoronavirus* |
| 6 | E-76 | *My. pilosus* | *Alphacoronavirus* |
| 7 | E-79 | *My. pilosus* | *unclassified Coronavirinae* |
| 8 | E-84 | *My. pilosus* | *Alphacoronavirus* |
| 9 | F-9 | *My. fimbriatus* | *unclassified Coronavirinae* |
| 10 | F-32 | *My. fimbriatus* | *unclassified Coronavirinae* |
| 11 | F-56 | *Rh. affinis* | *unclassified Coronavirinae* |
| 12 | F-58 | *Mi. schreibersii* | *Alphacoronavirus* |
| 13 | F-59 | *Mi. schreibersii* | *Alphacoronavirus* |
| 14 | F-87 | *Rh. sinicus* | *unclassified Coronavirinae* |
| 15 | H-22 | *Mi. schreibersii* | *unclassified Coronavirinae* |
| 16 | H-96 | *Mi. schreibersii* | *Alphacoronavirus* |

a. Bat ID indicates the information of the sampling site and the bat number.

**Table S13. Deposition of high-throughput sequencing data in open access database.**

| Sample accession | Sample ID | Counterpart bat individual | Accession (NMDC*) |
|---|---|---|---|
| Illumina sequencing | | | |
| NMDC20070104 | YN-pool 41 | H-15A | NMDC40041234 |
| NMDC20070105 | YN-pool 51 | C-45A | NMDC40041235 |
| NMDC20070106 | YN-pool 96 | H-75A | NMDC40041236 |
| NMDC20070107 | YN-pool 99 | C-40A | NMDC40041237 |
| NMDC20070108 | YN-pool 100 | D-16A | NMDC40041238 |
| NMDC20070109 | YN-pool 109 | F-92A | NMDC40041239 |
| NMDC20070110 | YN-pool 111 | A-92A | NMDC40041240 |
| NMDC20070111 | YN-pool 114 | H-54A | NMDC40041241 |
| NMDC20070112 | YN-pool 115 | H-82A and H-135A | NMDC40041242 |
| NMDC20070113 | YN-pool 117 | H-72A | NMDC40041243 |
| MGI sequencing | | | |
| NMDC20070114 | YN-P32-22 | C-22A | NMDC40041244 |
| NMDC20070115 | YN-P32-86 | C-86A | NMDC40041245 |
| NMDC20070116 | YN-P32-12 | C-12A | NMDC40041246 |
| NMDC20070117 | YN-P32-29 | C-29A | NMDC40041247 |
| NMDC20070118 | YN-P32-42 | C-42A | NMDC40041248 |
| NMDC20070119 | YN-P33-25 | C-25A | NMDC40041249 |
| NMDC20070120 | YN-P33-76 | C-76A | NMDC40041250 |
| NMDC20070121 | YN-P33-11 | C-11A | NMDC40041251 |
| NMDC20070122 | YN-P33-43 | C-43A | NMDC40041252 |

* NMDC, China National Microbiology Data Center (https://nmdc.cn/en)
